# Distinct Electrophysiological Signatures of Intentional and Unintentional Mind-Wandering Revealed by Low-Frequency EEG Markers

**DOI:** 10.1101/2023.03.21.533634

**Authors:** Adrien Martel, Nicolas Bruno, Ian H Robertson, Paul M Dockree, Jacobo D Sitt, Antoni Valero-Cabré

## Abstract

Mind-wandering is typically characterized by the common experience wherein attention veers off into thoughts unrelated to the task at hand. Recent research highlights the intentionality dimension of mind-wandering as a key predictor of adverse functional outcomes with intentional and unintentional task-unrelated thought (TUT) differentially linked to neural, behavioral, clinical, and functional correlates. We here aimed to elucidate the electrophysiological underpinnings of intentional and unintentional TUT by systematically examining the individual and collective discriminative power of a large set of EEG markers to distinguish between attentional states. Univariate and multivariate analyses were conducted on 54 predefined markers belonging to four conceptual families: ERP, spectral, information theory and connectivity measures, extracted from scalp EEG recordings prior to multidimensional reports of ongoing thought from participants performing a sustained attention task. We report here that on-task, intentional and unintentional TUT exhibit distinct electrophysiological signatures in the low frequency range. More specifically, increased features of the theta frequency range were found to be most discriminative between on-task and off-task states, while features within the alpha band were characteristic of intentional TUT when compared to unintentional TUT. This result is theoretically well aligned with contemporary accounts describing alpha activity as an index of internally oriented attention and a potential mechanism to shield internal processes from sensory input. Our study verifies the validity of the intentionality dimension of mind-wandering and represents a step forward towards real-time detection and mitigation of maladaptive mind-wandering.

## Introduction

Mind-wandering refers to the familiar experience wherein attention is engaged with thoughts that are uncoupled from current external stimuli or demands (Smallwood & Schooler, 2015). Estimated to occupy anywhere between 20% to 50% of waking mental activity (Killingsworth & Gilbert, 2010; Seli, Beaty, et al., 2018), mind-wandering has been found to predict a wide range of functional outcomes in both the laboratory and daily life (Mooneyham & Schooler, 2013). Although mind-wandering is known to purport benefits by promoting creativity, planning and problem solving (Chaieb et al., 2019), failure to appropriately suppress it has been linked to poor executive cognitive control (Unsworth & McMillan, 2014) and performance errors (Smallwood & Schooler, 2006) causing problems in educational (Smallwood et al., 2007), occupational (McVay et al., 2009) and operational settings (Baldwin et al., 2017). Given its ubiquity and link to a wide array of functional outcomes, understanding the nature of mind-wandering as a cognitive state, its neural basis and causal profile has emerged as a central goal within cognitive and clinical neuroscience (Christoff et al., 2016; Mittner et al., 2016; Seli, Kane, et al., 2018; Smallwood & Schooler, 2015).

While the definition of mind-wandering remains a matter of contention (Christoff et al., 2016; Seli, Kane, et al., 2018), the current work conceptualizes it as a hypernym for a phenomenologically diverse set of experiences, and as task-unrelated thought (TUT) when it occurs in the context of an explicit task. Mind-wandering is an intrinsically covert state with few, if any, overt behavioral markers, posing a unique set of challenges to scientific inquiry as well as real-time mitigation. To gain insight, researchers have leveraged the human capacity for introspection by asking participants to report on their experience. These ‘experience sampling’ approaches, range from the binary (i.e., asking participants to report being on- or off-task), to more granular approaches inquiring about multiple aspects of ongoing thought (for a review see (Weinstein, 2018)). The body of work that resulted from these approaches have exposed how mind-wandering experiences vary along several phenomenological and cognitive dimensions, such as metacognition (Christoff et al., 2009), emotional valence (Banks et al., 2016), or motivation (Robison et al., 2020).

A recent distinction with relevant practical and clinical applications concerns whether mind-wandering is engaged in with or without intention. While some have argued that TUTs primarily occur due to unintentional failures of executive control (McVay & Kane, 2009), prior work suggests that mind-wandering can, and does, occur with some intention (Seli et al., 2016). Intentionality refers to the degree to which mind-wandering results from a volitional reallocation of attention from the ongoing task towards TUT as opposed to mind-wandering ensuing from a dwindling of externally directed attention (El Haj et al., 2019; Grodsky & Giambra, 1990; Robison & Unsworth, 2018; Seli et al., 2013, 2014; Seli, Kane, et al., 2018; Seli, Maillet, et al., 2017; Seli et al., 2016; Seli, Smallwood, et al., 2015). The intentionality dimension of mind-wandering has recently emerged as a key factor with great explanatory power as supported by differential associations with specific content, neural, behavioral and clinical correlates (El Haj et al., 2019; Golchert et al., 2017; Martínez-Pérez et al., 2021; Seli, Ralph, et al., 2017; Seli et al., 2016; Seli, Smallwood, et al., 2015).

The “gold-standard” of experience sampling, online thought-probes, intermittently and unpredictably interrupt the task participants are engaged in and prompt them to classify their immediately preceding thoughts. Despite being the best method currently available to assess covert mental states and the advantages of providing an unmediated account of an individual’s attentional state, variability in individuals’ introspective ability as well as personal, contextual and motivational biases have raised doubts over its validity (Konishi & Smallwood, 2016; Seli, Jonker, et al., 2015; Weinstein et al., 2018). Additional issues relate to the modest-to-weak correlations between TUT rates in laboratory tasks and daily-life (Kane et al., 2017; McVay et al., 2009) and the widespread use of dichotomous thought-probes (Weinstein, 2018), i.e., simply contrasting on- and off-task, which have been found to inflate off-task reports (Seli, Beaty, et al., 2018). Several approaches have been attempted to remedy these limitations. Researchers have worked on refining subjective measures by leveraging the heightened metacognitive ability of expert meditators (Ellamil et al., 2016), by expanding thought-probes categories to include more dimensions (e.g. Robison et al., 2020) or confidence ratings (Seli, Jonker, et al., 2015), by testing the construct validity of experience sampling (Kane et al., 2021) or by identifying the optimal thought-probe frequency (Welhaf et al., 2022). Nevertheless, thought sampling requires interruptions of the task, which can increase participants’ meta-awareness about the content of their thoughts (Zedelius et al., 2015) and alters TUT reports and rates (Seli et al., 2013). A novel approach has hence been proposed that aims at curbing the reliance on thought sampling by corroborating self-reports with objective measures, e.g., errors and reduction in response variability (Smallwood & Schooler, 2015) via a “triangulation” process between several markers of TUTs (Konishi & Smallwood, 2016). Based on the assumption that objective measures are ideal markers of cognitive processes (Andrews-Hanna et al., 2018), numerous studies have investigated the behavioral and physiological correlates of mind-wandering. Previous work identified behavioral markers of mind-wandering, e.g., response time (McVay & Kane, 2009)), task-related measures such as driving performance (Baldwin et al., 2017), and physiological measures, including skin conductance (Smallwood et al., 2004) and pupillometry (Groot et al., 2021). Although these findings show promise for objective measures to complement or replace self-reports, these markers remain heavily reliant on specific tasks or contexts, require specialized equipment or setups, and mostly fail to inform on the brain dynamics underlying TUTs. In comparison, neural measures provide a direct window into the dynamics of the neurocognitive processes underlying covert mental states such as mind-wandering. Based on robust markers, the exquisite temporal resolution inherent to electromagnetic measures of brain activity allows for the real-time detection of mental states, opening avenues for the online mitigation of adverse outcomes linked to mind-wandering (Gouraud et al., 2021) and ultimately, the obviation of subjective methods.

### Electrophysiological markers of mind wandering

Identifying electrophysiological correlates of mind-wandering has long been a goal within the field with early scalp EEG studies mainly investigating event-related potentials (ERPs). Early potentials such as P1 and N1, which reflect evoked responses to sensory stimuli, have been found to decrease when participants reported being engaged in TUT (Baird et al., 2014; Dong et al., 2021; Gouraud et al., 2021; Julia W. Y. Kam et al., 2010; Martel et al., 2019). The amplitude of the P3, which indexes the general level of cognitive processing or allocation of attentional resources (Polich, 2007), has also been observed to be reduced during TUTs in various studies (Baldwin et al., 2017; Dias da Silva et al., 2022; Gouraud et al., 2021; Groot et al., 2021; Kam et al., 2010; Smallwood et al., 2008). In line with the perceptual decoupling hypothesis of mind-wandering (Smallwood & Schooler, 2015), these findings suggest a strong link with a general attenuation in cortical processing of external stimuli causing a drop in short-term performance (Mooneyham & Schooler, 2013; Randall et al., 2014) and overall vigilance (Braboszcz & Delorme, 2011).

In addition to ERP activity, multiple studies have investigated the relationship between mind-wandering and EEG oscillatory activity. Analyses of the time-frequency decomposed signal within canonical frequency bands (i.e., delta (1-4Hz), theta (4-8Hz), alpha (8-14Hz), beta (15-30Hz), and gamma (30-50Hz)) have yielded even more variable results compared to ERP studies (for a review see Kam et al., 2022). Alpha oscillations centered at 10Hz, are the most prominent rhythm in the healthy brain and have long been associated with mental states cognate to mind-wandering. For example, while alpha-band activity is reduced in response to increased attentiveness and perceptual stimulation (Thut et al., 2006), it is increased during waking eyes-closed compared to eyes-open conditions (Adrian & Matthews, 1934), during rest (Compton et al., 2011), and during attentional lapses (Macdonald et al., 2011; Martel et al., 2014; O’Connell et al., 2009). Greater alpha-band activity has also been linked with internally oriented attention (Benedek et al., 2014; Ceh et al., 2020; Hanslmayr et al., 2011), mental imagery (N. R. Cooper et al., 2003) and activity in the default mode network (Knyazev et al., 2011).

Overall, there is substantial evidence for the key role alpha oscillations play in gating information flow throughout the brain (Jensen & Mazaheri, 2010; Klimesch, 2012), and as a top-down inhibitory control mechanism to maintain task performance by suppressing task-irrelevant information (Händel et al., 2011). Numerous studies have reported greater alpha power over frontal, central, parietal and occipital scalp areas during periods of TUT (Arnau et al., 2020; Baldwin et al., 2017; Ceh et al., 2020; Compton et al., 2019; Dias da Silva et al., 2022; Groot et al., 2021; Hanslmayr et al., 2011; Jin et al., 2019; Macdonald et al., 2011; Martel et al., 2019), positioning it as a promising EEG signature of mind-wandering despite some evidence to the opposite (Baird et al., 2014; Braboszcz & Delorme, 2011).

Together with the attenuation in ERP amplitudes, greater alpha-band activity suggests a drop in the cortical processing of the external environment as attention is redirected internally during TUT. This is consistent with the executive function model of mind-wandering (Smallwood & Schooler, 2006, 2015), which argues that executive resources need to be decoupled from sensory input during mind-wandering to shield internal processes from interferences and permit TUTs to unfold uninterrupted (Smallwood, 2013). Thus far, only two studies have examined EEG differences between intentional and unintentional TUTs. These found ERPs and in particular the P3 component to be diminished during off-task states when compared to on-task while greater alpha activity was found to be linked with more intentional forms of TUT (Kam et al., 2021; Martel et al., 2019).

While increased alpha oscillations during TUT is a relatively consistent finding, patterns of activity in the theta band have shown more variability, with some studies observing greater theta activity during TUTs compared to on-task states (Arnau et al., 2020; Martel et al., 2019; Polychroni et al., 2022; van Son, De Blasio, et al., 2019) and others reporting the opposite pattern (Kirschner et al., 2012; Wamsley & Summer, 2020). Ample evidence suggests that theta rhythms within and across brain regions subserve executive control (Cavanagh & Frank, 2014) and attentional functions (Helfrich et al., 2018) with some studies reporting increased frontal theta power when participants performed tasks imposing demands on externally oriented attention (Clayton et al., 2015; Kubota et al., 2001).

### Detection of task-unrelated thoughts

Altogether, findings from electrophysiological studies point towards a reliable EEG signature for TUTs which can be readily exploited with machine learning techniques. Accordingly, recent research demonstrates the possibility of predicting the occurrence of TUTs based on EEG measures (Chen et al., 2020; Dhindsa et al., 2019; Dong et al., 2021; Jin et al., 2019; Polychroni et al., 2022). Different classification approaches have been utilized on varied features of EEG and despite some variations, certain features appear most characteristic of TUTs, e.g., P3 and alpha (Dong et al., 2021; Groot et al., 2021; Polychroni et al., 2022), corroborating electrophysiological studies.

In this study, we aim at disentangling the EEG signatures of intentional and unintentional TUTs and determine whether they differ from on- and off-task. To this end, we assessed the overall discriminative ability of a wide range of putative EEG markers by performing a large-scale analysis on a range of predefined EEG features and testing their individual and collective ability to discriminate between both on- and off-task activity, as well as iTUT and uTUT.

The approach used has been developed and successfully employed to robustly extract electrophysiological markers of consciousness across contexts and protocols identifying features in the alpha and theta band as indexes of consciousness (Engemann et al., 2018; Sitt et al., 2014). Given that mind-wandering states are closely linked to conscious content (Schooler, 2002; Smallwood & Schooler, 2015) and to conscious access (Dias da Silva et al., 2022), we are extending this feature-extraction tool to TUT. Applying this approach to an EEG dataset recorded during a standard test of sustained attention represents the first attempt at systematically investigating the discriminative features of intentional and unintentional TUT. As such, we expected to identify ERP markers, e.g., P3, as most discriminative between on- and off-task states, and for markers in the alpha band to distinguish intentional from unintentional TUT.

## Materials & Methods

### Dataset

Twenty-six participants (12 females, age M: 25 SD: 4.3) with normal or corrected-to-normal vision and no history of neurological or psychiatric disease, volunteered or received partial course credits to participate in the study. All procedures were approved by Trinity College Dublin ethics committee and conducted with adherence to the Declaration of Helsinki. Participants were informed extensively about the experiment, and all gave written consent.

### Task stimuli and paradigm

Participants were seated in a soundproof, electrically shielded, dimly lit room and performed a fixed version of the sustained attention to response task (SART (Robertson et al., 1997)). The fixed SART used here is a computerized go/no-go task requiring participants to withhold behavioral response to infrequent no-go targets (no-go: 6) presented amongst a background of frequent and sequentially presented non-targets (go: 1 to 5 and 7 to 9). A monitor at a viewing distance of approximately 70 cm, sequentially and centrally displayed digits from 1 to 9 for 250 ms with an inter-stimulus interval (ISI) of 2316.5 ms (see Fig. 1). This ISI optimizes for the tradeoff between inducing a maximum number of mind-wandering episodes while not exacerbating the task’s difficulty by being too monotonous and imposing excessive demands on attentional resources. Stimuli were presented at a font size of 140 in Arial font using the Presentation software package v19.0 (www.neurobs.com). Participants were instructed to button press as fast as possible to the go digits and lock their response to the offset of the stimulus, a response strategy that has been successfully applied to minimize both the inter-individual variability in response times and speed-accuracy tradeoffs (O’Connell et al., 2009). The SART was composed of three blocks of a duration for a minimum of 8 min and a maximum of 15 min, thus, the total running time of the task ranged from 24 min to 45 min. The task was composed of between 800 and 1200 trials approximately. The duration of each SART block was relative to participants’ answers to the self-reports with each block lasting for at least 8 min or until three reports for each of the main categories of interest were given (on-task, TUTs). The task took on average 38.7 min (*SD*: 4.8) with an average of 65.3 thought probes (*SD*: 31.2) of which 71.4 % were of the self-caught variant.

**Figure 1.**
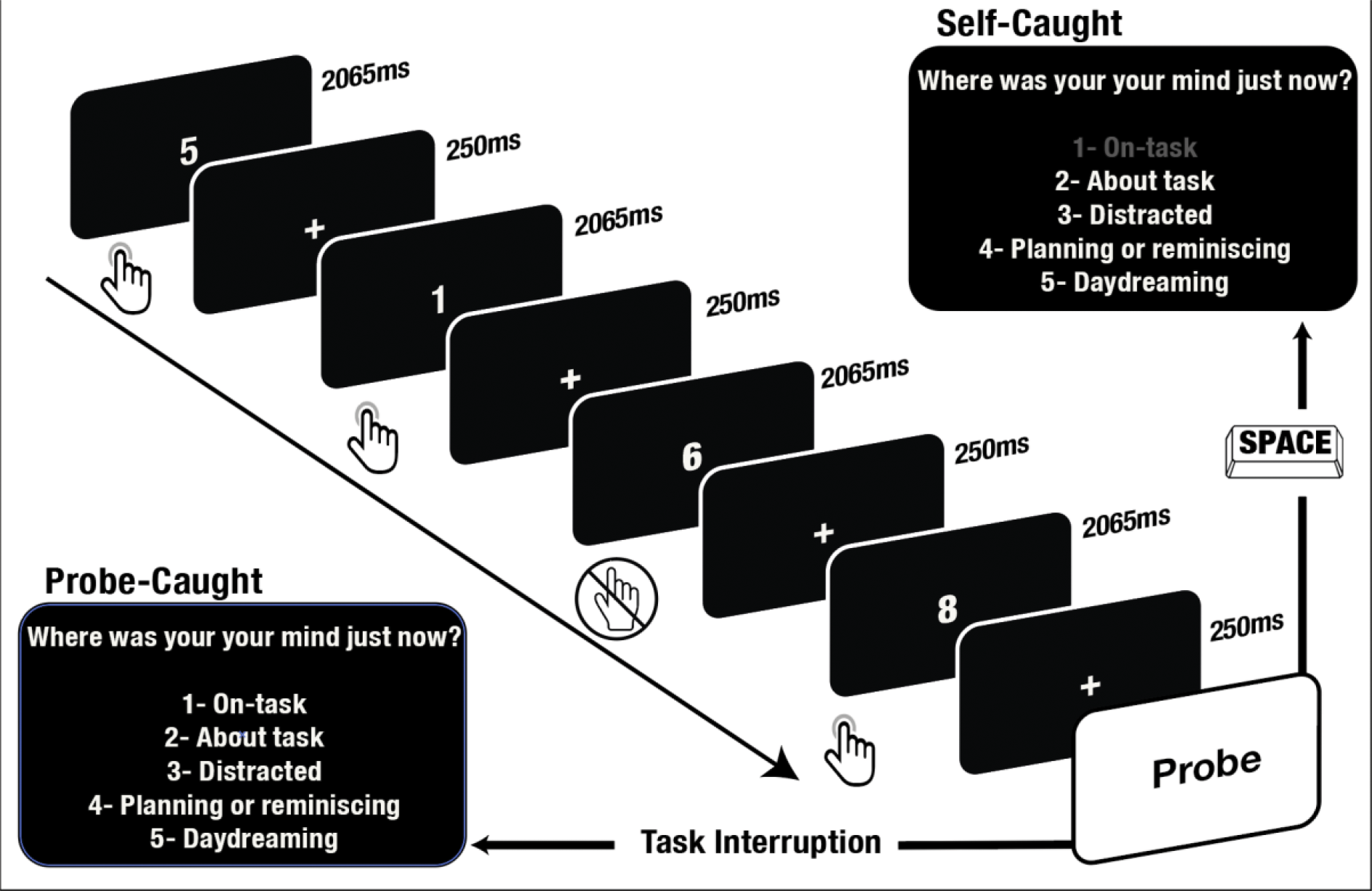
Sustained-Attention-to-Response Task (SART) featuring thought probes. Participants observed a continuous sequence of single digits, pressing a button for each digit except the number 6 (targets). Attentional state was assessed intermittently through probe-caught probes (occurring on average every 30 trials) or self-caught probes initiated by pressing the space bar.

Throughout the SART participants were prompted to report on their attentional state in two different ways: Probe-caught and self-caught thought probes. Probe-caught probes would intermittently and pseudo-randomly interrupt the task every 12, 18, 24, 30, 36, 42, or 48 trials, on average every 30 trials (∼ 28s, 42s, 56s, 76s, 83s, 97s, 111s, on average every 76s). Upon interruption participants were asked the question “Where was your mind just now?” and prompt them to classify their ongoing thoughts prior to the interruption according to five categories:

1. on task (focused attention)
2. about the task (task-related thoughts)
3. distracted (internal or external interference)
4. n future plans or memories (intentional TUT)
5. daydreaming (unintentional TUT)

In parallel, participants were instructed to trigger a self-caught probe by pressing the space bar whenever they realized that they were no longer on-task. While participants were instructed to only report being on-task during probe-caught reports, the ‘on-task’ option remained available during self-caught probes to account for accidental button presses. Participants were trained in the correct categorization of attentional states prior to the start of the experiment, with different examples for each category and a quiz to test their understanding. For convenience and to avoid confusion, the categories for intentional and unintentional TUT were named ‘on future plans or memories’ and ‘daydreaming’, respectively. To isolate TUT from other forms of mind-wandering, participants who found themselves having task-related thoughts, e.g., thoughts about their response strategy, were instructed to choose the ‘about task’ category (Stawarczyk et al., 2013). Similarly, when thoughts were related to internal sensations or external distractions, e.g., an itch or a noise in the environment, participants were instructed to report the ‘distracted’ category.

### EEG acquisition and preprocessing

The EEG was recorded from 64 active channels placed on a cap according to the international 10-20 reference system, using the BioSemi ActiveTwo system (www.biosemi.com). Continuous EEG data were amplified and digitized at 512 Hz, and bandpass filtered between 0.5 and 45 Hz. To assess eye movements and blinks, 4 electrooculography channels (EOGs) were used; two placed above and beneath the left eye and one on the outside of each eye. The preprocessing of EEG was performed with the MNE-python software package(Gramfort et al., 2014) (www.mne.tools). EEG data was down-sampled to 250 Hz before being high-pass filtered at 0.5 Hz and low-pass filtered at 45 Hz. Channels with excessively noisy signals were removed. Following Sitt et al.(Sitt et al., 2014), the EEG data was segmented into epochs spanning -200 ms to 600 ms relative to SART stimuli and baseline corrected with respect to the pre-stimulus period (−200 to 0 ms). Noisy epochs were removed automatically with the *Autoreject* package (https://autoreject.github.io/). To correct for ocular and muscle artifacts, an ICA decomposition with the FastICA method was performed(Hyvarinen, 1999) and artifactual components were removed. Previously removed channels were interpolated using a spherical spline interpolation before re-referencing using a common average reference. Additionally, the ERP component of all epochs was subtracted for the computation of all non-evoked markers. From this point on, only the 5 epochs prior to a report (∼10 s) were considered and labelled according to the category of thought reported.

### Analysis

Univariate and multivariate pattern analyses were conducted over all 54 markers across two contrasts. The first contrast compared probe-caught on- and off-task conditions with the off-task condition consisting of combined intentional and unintentional TUT epochs to balance sample size (see Martel et al., 2019). The second contrast consisted of iTUT and uTUT reports from self-caught probes. For the univariate and multivariate analyses, we computed a total set of 27 markers drawn from Sitt et al. (2014) and Engemann et al. (2018) with each belonging to one of four conceptual families of markers: event-related potentials (ERPs), Spectral, Information Theory, or Connectivity (see Table 1). Although Sitt et al. (2014) employed a larger set of markers we used a subset since many were found to be redundant and/or yielding poor performance. The set of markers used here mirrors the selection by Engemann et al.(2018) with the addition of information theory and connectivity markers for all spectral bands, and excluding markers specific to their experimental methodology. For a detailed description and discussion of the markers see Sitt et al. (2014). All markers were computed using the NICE library (available at (https://github.com/nice-tools/nice) for each of the 5 epochs immediately preceding a thought probe and linked to the condition corresponding to the self-report category. This time window of approximately 12 sec preceding thought-probes is broadly consistent with previous analyses (Baird et al., 2014; Baldwin et al., 2017; Braboszcz & Delorme, 2011).

**Table 1.** Description of the full list of EEG-markers used and the category to which they pertained.

| Acronym | Marker | Category |
| --- | --- | --- |
| CNV | Contingent Negative Variation | ERP |
| P1 | P100 evoked potential | ERP |
| P3a | P3a evoked potential | ERP |
| P3b | P3b evoked potential | ERP |
| $\alpha$ | Alpha PSD | Spectral |
| $\vert \alpha\vert $ | Normalized Alpha PSD | Spectral |
| $\beta$ | Beta PSD | Spectral |
| $\vert \beta\vert $ | Normalized Beta PSD | Spectral |
| $\delta$ | Delta PSD | Spectral |
| $\vert \delta\vert $ | Normalized Delta PSD | Spectral |
| $\gamma$ | Gamma PSD | Spectral |
| $\vert \gamma\vert $ | Normalized Gamma PSD | Spectral |
| $\theta$ | Theta PSD | Spectral |
| $\vert \theta\vert $ | Normalized Theta PSD | Spectral |
| MSF | Median Power Frequency | Spectral |
| SE90 | Spectral Edge 90 | Spectral |
| SE95 | Spectral Edge 95 | Spectral |
| SE | Spectral Entropy | Spectral |
| K | Kolgomorov Complexity | Information Theory |
| PE $\alpha$ | Permutation Entropy Alpha | Information Theory |
| PE $\beta$ | Permutation Entropy Beta | Information Theory |
| PE $\delta$ | Permutation Entropy Delta | Information Theory |
| PE $\gamma$ | Permutation Entropy Gamma | Information Theory |
| PE $\theta$ | Permutation Entropy Theta | Information Theory |
| wSMI $\alpha$ | weighted Symbolic Mutual Information Alpha | Connectivity |
| wSMI $\beta$ | weighted Symbolic Mutual Information Beta | Connectivity |
| wSMI $\delta$ | weighted Symbolic Mutual Information Delta | Connectivity |
| wSMI $\gamma$ | weighted Symbolic Mutual Information Gamma | Connectivity |
| wSMI $\theta$ | weighted Symbolic Mutual Information Theta | Connectivity |

The computation of each marker yielded multiple observations per channel, per epoch, per time point, and/or per frequency bins depending on the family of the marker. These observation points were aggregated first in the time/frequency/sensor domain depending on marker type, before being averaged across all sensors. Finally, the last 5 epochs corresponding to one condition were aggregated via an average and the standard deviation, the latter being a measure of the variations between epochs (see Fig. 2). This yielded two different measures per epoch and marker, giving a total of 54 markers (27 average, and 27 variation measures). The markers were labeled according to their type and to the final processing step, i.e., ‘mean’ or ‘std’ (e.g., *α*_*mean*_or *α*_*std*_). All analysis scripts are publicly available in https://github.com/Nicobruno92/mw_markers_project.

**Figure 2.**
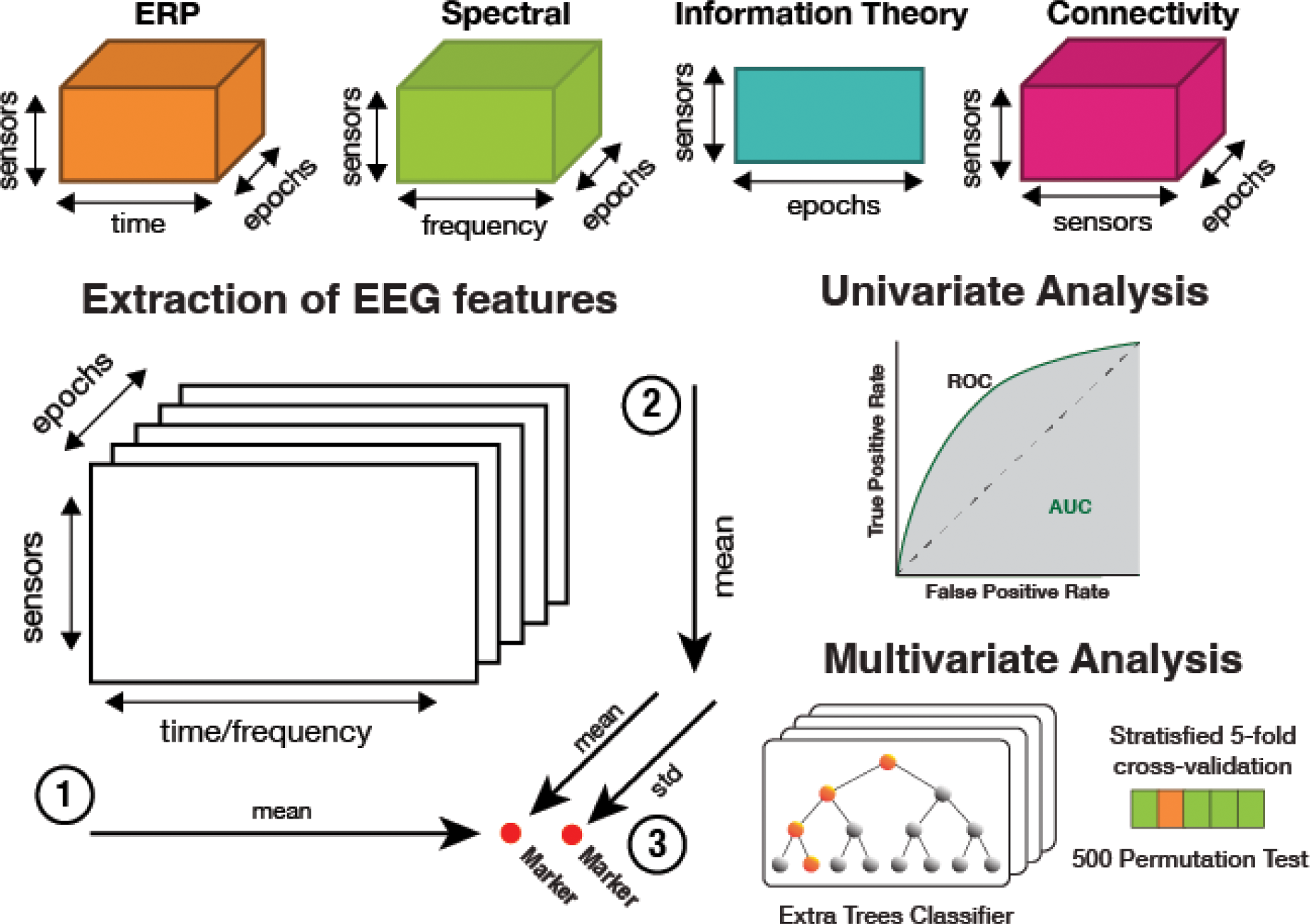
EEG feature extraction pipeline for univariate and multivariate analyses. EEG markers were categorized into four conceptual families: evoked responses, spectral, information theory, and connectivity. These were aggregated in three steps for the five SART trials preceding a probe response. (1) EEG features from ERP, spectral, or connectivity families were averaged across time, frequency, or sensor dimensions, respectively. (2) Markers were averaged over all EEG channels to yield one unique value per epoch. (3) Two features were extracted from each marker (indicated by red dots) by calculating both mean and standard deviation. Markers were labeled according to type and final computation step, e.g., standard deviation of alpha oscillations was labeled ɑ_std_. Univariate analysis employed ROC-AUC metric for each of the 54 markers to assess individual classification performance for both contrasts (on-task/off-task and iTUT/uTUT). Multivariate analysis evaluated the collective discriminative ability of all markers for both contrasts using Extra Trees Classifier.

### Statistical analysis

#### Univariate analysis

The univariate analysis of individual markers followed the same procedure as Sitt et al.(2014). The area under the curve (AUC) of the receiving operating characteristic (ROC) curve served as an estimate of the discriminative ability for a given marker. The ROC curve depicts the false-positive rates (FPR) against the true positive rates (TPR), with each point of the curve representing an FPR/TPR pair for varying decision thresholds. A decision threshold with perfect discrimination would yield an FPR of 0 and a TPR of 1. The result of computing all possible decision thresholds, is the area under the curve (AUC) of the ROC as a metric of performance. Any AUC value between 0.5 and 1 indicates positive discriminative ability of a given feature for the first category (on-task or iTUT) while AUC values between 0 and 0.5 denotes the reverse pattern, namely positive discriminative ability of a given feature for the second category (off-task or uTUT). And an AUC of 0.5 indicates chance levels of discrimination. To assess the significance of the discrimination for each marker the Mann-Whitney U test for independent samples was computed. These analyses were applied for the entire set of 54 markers previously described, composed of 27 averages and 27 standard deviations. Given the number of markers, statistical significance was corrected for multiple comparisons using the false discovery rate (FDR) method.

#### Multivariate pattern analysis (MVPA)

In keeping with the methods applied in Engemann et al. (2018), we used an *Extra-trees* classifier (Geurts et al., 2006) from the Scikit-learn python library (Pedregosa, 2011) for the MVPA. Extra-Trees classifiers are non-parametric models that perform robustly and are less sensitive to the scale of the input data (Engemann et al., 2018). This category of model is also more efficient at handling the type of dataset we obtained, called ‘wide’ because it contains more variables than observations. A further advantage of this type of model is the possibility of outputting feature importance scores which provide additional information on the discriminative capacity of individual features. In addition to the univariate correlation, this score also considers the interdependency with other variables. For the classifier, we used 1000 trees with entropy as impurity criterion and the other parameters set to default. The multivariate model was trained and tested based on the same 54 markers used during the univariate analyses.

For cross-validation, Monte-Carlo cross-validation was used with the training set size of 80 % and a testing set of 20 % with 5 iterations. To test the statistical significance of our model, we applied a 1000-permutation test for the same cross-validation procedure, yielding 1000 samples with shuffled labels which were compared to the real cross-validation sample.

## Results

Considered in the analysis were 306 probe-caught and 935 self-caught reports, of which 94 were categorized as on-task, 82 as off-task, 286 as iTUT and 250 as uTUT (see Table 2 for details).

**Table 2.** Count of self-reports per thought category, separated by probe-caught and self-caught probes.

| Probe | On-task | About-task | Distraction | iTUT | uTUT | Total |
| --- | --- | --- | --- | --- | --- | --- |
| Probe-caught | 94 | 82 | 48 | 39 | 43 | 306 |
| Self-caught | - | 200 | 199 | 286 | 250 | 935 |

### On- vs off-task

#### Univariate analysis of on-/off-task

The univariate analysis revealed that the most discriminative marker for the on-/off-task contrast was the normalized power of the theta frequency band, with both the average (|θ|_mean_: *AUC =* 0.612, *p*_*uncorrected*_ = 0.01) and the standard deviation (|θ|_std_: *AUC* = 0.600, *p*_*uncorrected*_ = 0.023) reaching statistical significance (see Fig. 3A). Although effect sizes showed discriminative values (i.e., AUC values above/below 0.50), none of the measures for this contrast survived FDR correction.

**Figure 3.**
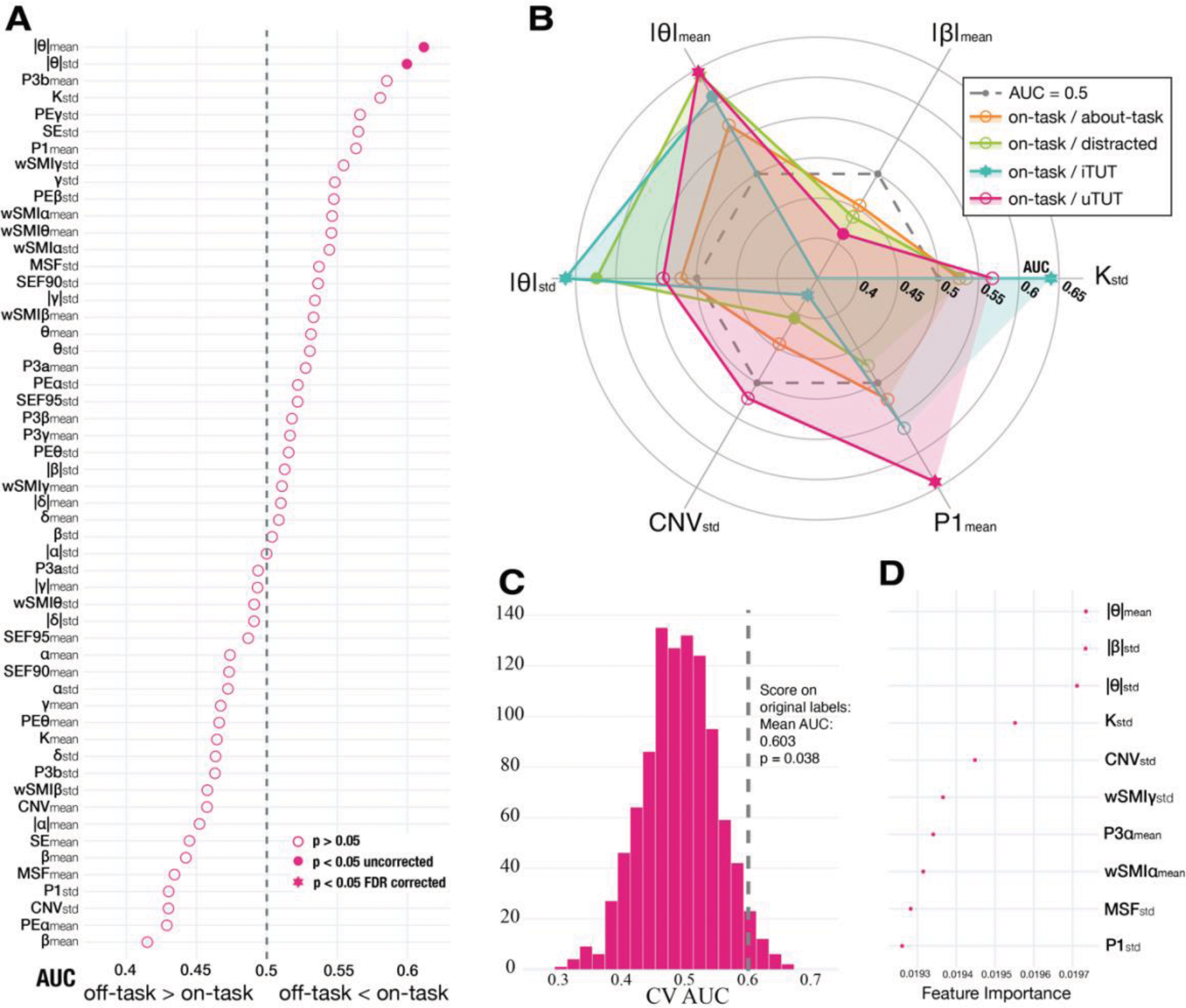
Discrimination measures for EEG markers using AUC. AUC>0.5 indicates higher measure for on-task than the other condition; AUC<0.5 indicates the opposite pattern; AUC=0.5 implies no discrimination. Filled circles represent *p*<0.05 in a Mann-Whitney U test before multiple comparison correction; filled stars indicate significance after FDR correction. (A) AUC values of all markers for on-/off-task contrast, ranked by significant markers not surviving multiple comparison correction (filled circles) and non-significant markers (empty circles) in decreasing AUC order. (B) Scatter polar plot of AUC values for markers significant after correction (filled stars) in at least one comparison with on-task condition (about-task, distracted, iTUT, or uTUT). (C) Histogram of permutation tests evaluating statistical significance of MVPA for on-/off-task model, with histogram bins representing AUC of 1000 permuted models with shuffled labels; dashed line shows mean AUC for cross-validated models with correct labels. (D) Average feature importance measures from MVPA for top 10 features in on-/off-task model.

#### Multivariate Pattern Analysis (MVPA) of on-/off-task

To assess the collective discriminative ability of the markers for the on-/off-task contrast, an Extra trees classifier was trained on all the markers included in our prior univariate analysis. The accuracy of the classifier was computed via the mean of the 5-fold cross-validation. The classifier achieved above-chance performance with an *AUC M* = 0.603. The statistical significance of this model was determined using a 1000 permutations test revealing that the accuracy obtained was significantly above chance level *p* = 0.042 (See Fig. 3C). Moreover, we obtained the feature importance of this classifier as another measure of univariate discriminative ability for each marker (see Fig. 3D). Corroborating the results of our univariate analysis, theta mean, and theta standard deviation were amongst the most important features, with the second most important feature being the average of beta band normalized power (see Suppl. Fig. 2).

#### Contrasting on-task with the other conditions

To further explore whether the significant markers for on-/off-task were specific to this comparison or resulted from idiosyncrasies of the on-task condition, we contrasted on-task against the other four categories of thought (i.e., about-task, distracted, iTUT, and uTUT). To balance the samples across classes we used the Synthetic Minority Over-Sampling Technique (SMOTE) which generates synthetic values by linear interpolations of nearest-neighbor values. This approach has been previously used for TUT classification given that mind-wandering experiments typically have unbalanced samples (Dong et al., 2021). Subsequently, we repeated the same univariate procedure described in the previous section for each of the contrasts (on-task/about-task, on-task/distracted, on-task /iTUT, and on-task/uTUT).

This analysis yielded normalized theta power as the only significant marker against all unfocused states conditions (distracted, iTUT and uTUT; see Fig. 3B) and was consistently higher for on-task than for any other category of thought. Moreover, the variation measure of this marker reached significance after FDR correction for the contrasts of on-task against distracted and iTUT. Additionally, normalized beta power was found to be higher for all the unfocused states (distracted, iTUT and uTUT) compared to on-task but was only significant for uTUT. It is also interesting to remark that ERP component P1 was found to be significant after multiple comparison correction and higher for on-task when contrasted with uTUT. The one contrast for which no markers were significant, was between on- and about-task, most likely because of the similarity between these two categories.

### Intentional vs Unintentional TUT

#### Univariate Analysis of iTUT/uTUT

The univariate analysis of the iTUT/uTUT contrast revealed permutation entropy in the theta frequency range averaged across trials (PEθ_mean_: *AUC* = 0.625, *p*_*FDR*_ < 0.001) as the most discriminative marker, with higher values for iTUT when compared to uTUT (see Fig. 4A). Next in order, both the normalized (|α|_mean_: *AUC* = 0.593, *p*_*FDR*_ = 0.002) and non-normalized (α_mean_: *AUC* = .598, *p*_*FDR*_ = .004) mean alpha power were found to be significantly higher for the iTUT condition than for uTUT. The most discriminative marker characterizing uTUT when compared to iTUT that was increased normalized delta power (|δ|_mean_: *AUC* = 0.411, *p*_*FDR*_ = 0.005). Additionally, unnormalized beta power (β_mean_: *AUC* = 0.576, *p*_*FDR*_ = 0.026) and permutation entropy (PE β_mean_: *AUC* = 0.431, *p*_*corrected*_ = 0.006) were also found to be significant markers after FDR correction for the classification between iTUT and uTUT. For more details on this analysis see Supplementary Table 2.

**Figure 4.**
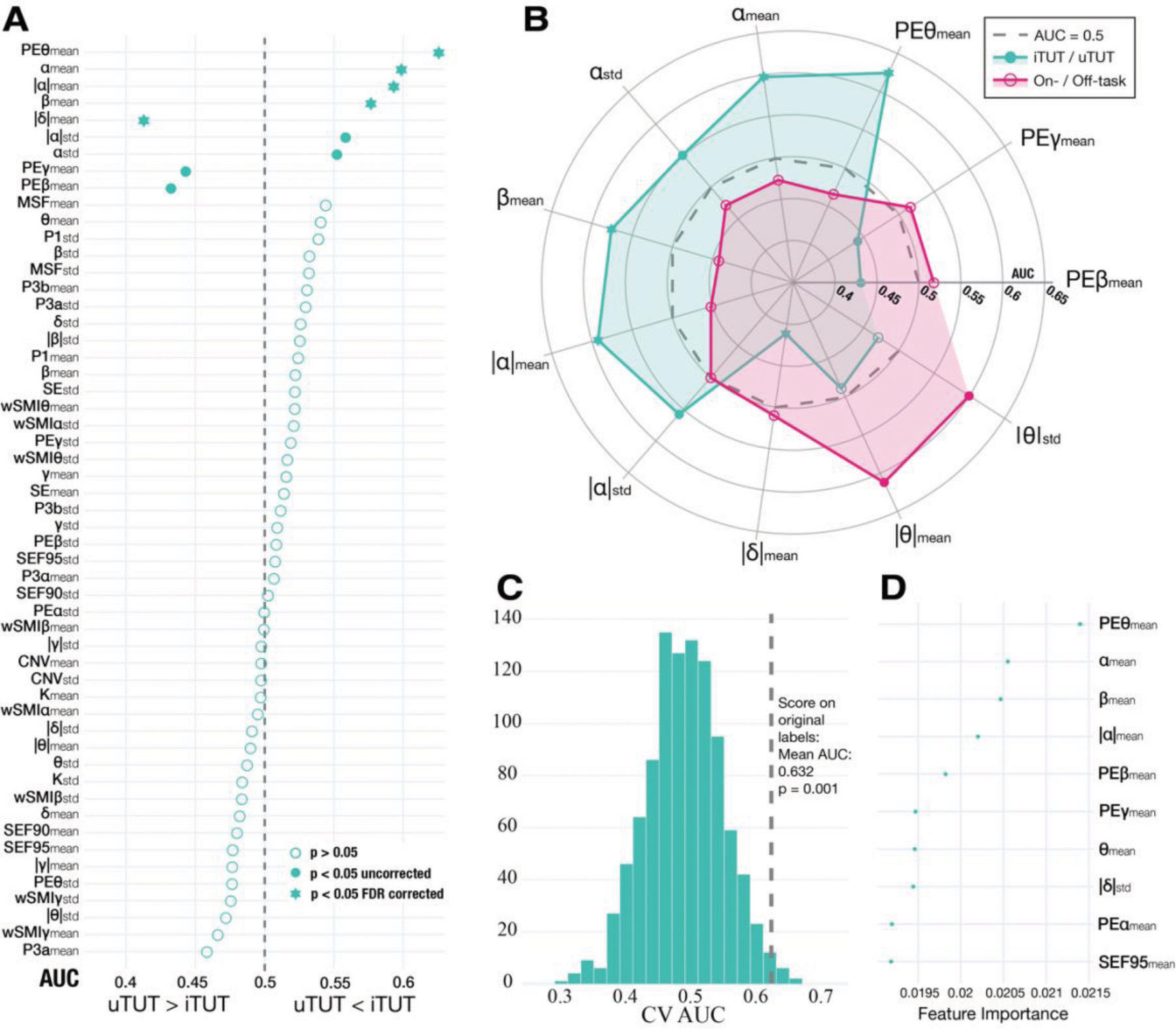
Discrimination measures for iTUT/uTUT and on/off-task contrasts using AUC. AUC>0.5 indicates higher measure for the first condition (iTUT or on-task) than the second (uTUT or off-task); AUC<0.5 indicates the reverse pattern; AUC=0.5 implies no discrimination. Filled circles represent *p*<0.05 in a Mann-Whitney U test before multiple comparison correction; filled stars indicate significance after FDR correction. (A) AUC values of all markers for iTUT/uTUT comparison, ranked by significance after multiple comparison correction (filled stars), significant markers not surviving multiple comparison correction (filled circles), and non-significant markers (empty circles). (B) Scatter polar plot of AUC values for significant markers in on-/off-task or iTUT/uTUT contrasts. (C) Histogram of permutation tests evaluating statistical significance of MVPA for iTUT/uTUT model, with histogram bins representing AUC of 1000 models with shuffled labels; dashed line shows mean AUC for cross-validated models with correct labels. (D) Average feature importance measures from MVPA for top 10 features in iTUT/uTUT model.

The same univariate analysis was repeated to test if there was any marker able to separate the three unfocused categories (i.e., distracted, iTUT and uTUT). Our analyses revealed that none of the significant markers were shared across the three contrasts (see Supplementary Fig. 3). However, most of the markers that were found to be significant for the contrast between iTUT and uTUT, were also significant for the iTUT and the distracted condition (e.g., PE θ_mean_, PE β_mean_, PE ɣ_mean_, |α|_mean_, α_mean_, |δ|_mean_ and β_mean_). Moreover, no marker was significant for the contrast of uTUT and the distracted condition after FDR correction.

#### Comparison of univariate results for on/off-task and iTUT/uTUT

Finally, to examine whether similarities exist in discriminative markers across both contrasts, we mapped the two sets of significant measures the on-/off-task and the iTUT/uTUT contrast, in a polar plot (Fig. 4B). The resulting figure shows two distinct constellations of significant markers, suggesting that iTUT and uTUT are characterized by unique patterns of EEG markers.

Furthermore, we assessed whether the computed average and standard deviation features were equally informative in terms of their classification power across the two contrasts (on-/off-task and iTUT/uTUT). For this analysis, we mirrored the AUC values for all the markers (i.e., computing the difference with 1 for each of the *AUC* < 0.5; see Suppl. Fig. 1) to obtain an absolute value of the classifications and avoid direction bias. Then, to determine whether AUC distributions were statistically different, we computed a Mann Whitney U test between the mirrored AUC values of each marker for each contrast (i.e., the AUC of the on-/off-task contrast against the AUC values for the iTUT/uTUT contrast) for the averaged features. This was also performed separately for the standard deviation features (see Suppl. Fig. 1). While this analysis yielded no significant differences for the average markers (*U* = 248; *p* = 0.362), significant differences were found when this was repeated over standard deviations (*U* = 169; p = 0.018). In the latter case, standard deviations were more discriminative for the on-/off-task comparison (*Median AUC* = 0.536) than for the iTUT/uTUT contrast (*Median AUC* = 0.517).

#### MVPA of iTUT/uTUT

The MVPA for the iTUT/uTUT contrast revealed a modest increase in classification accuracy in comparison with the previous analysis, with an *AUC M* = 0.632 for the cross-validation results. Further, the permutation test determined the model classification performance to be higher than chance, *p* = 0.002 (see Fig. 4C). Moreover, the feature importance from this model was computed, as an added indicator of the discrimination of each of the measures. The results proved similar to those obtained with our univariate analysis; permutation entropy in the theta frequency band emerged as the most important feature, followed by beta and alpha power (see Fig. 4D).

## Discussion

Amongst the confluence of factors that contributed to mind-wandering becoming a research focus, its prevalence as a mental activity and robust association with adverse functional outcomes have been particularly catalytic. The intrinsically covert nature of mind-wandering constitutes a central challenge to scientific inquiry and the main driver for the widespread adoption of experience sampling as the default method of collection. However, subjective reports are prone to biases prompting a growing number of studies to identify objective measures by, for example, developing machine learning models that can reliably detect attentional states based on EEG. Concurrently, recent views from both, psychology(Seli, Kane, et al., 2018) and neuroscience(Wang et al., 2018) have established mind-wandering as a multifaceted construct that varies along several cognitive dimensions of which intentionality has emerged as a key predictor of functional outcomes(Julia W. Y. Kam et al., 2022). These circumstances call for more fine-grained self-reports incorporating the intentionality dimension to accurately study the complex mechanisms underlying off-task thought and help unravel the inconsistencies in electrophysiological correlates of TUT reported across EEG studies, in particular regarding oscillatory markers (Kam et al., 2022).

Using the SART and EEG, our study aimed at identifying which, from a set of 52 predefined markers belonging to one of four families of EEG measures (ERP, spectral, connectivity, permutation entropy), were most characteristic of on- and off-task states, and of intentional and unintentional task-unrelated thoughts (TUTs), respectively. Both our univariate and multivariate analysis demonstrated that on-task states are reliably characterized by greater normalized power and variance in theta range than off-task states, and that intentional TUT (iTUT) were characterized by increased theta permutation entropy and greater alpha power measures when compared to unintentional TUT (uTUT). More specifically, the most discriminative EEG marker for on-task states was normalized power in the theta frequency range (4-7 Hz) an outcome that was further corroborated by an MVPA feature importance analysis and by contrasting the on-task condition with the other categories of thought sampled (i.e., about-task, distracted, iTUT, uTUT). The same analyses performed on the iTUT/uTUT contrast revealed that iTUT was associated to greater permutation entropy at the theta frequency range and increased alpha spectral measures. Also, there was no overlap in discriminative markers across contrasts, suggesting that on-task, iTUT and uTUT have distinct electrophysiological signatures. This is consistent with a growing body of literature showing that intentional and unintentional mind-wandering rely on separate cortical architectures with specific patterns of neural activity(Golchert et al., 2017; Martel et al., 2019; Seli et al., 2016).

One of the principal results of this study was that features of alpha activity did not differentiate on-from off-task states but did significantly discriminate between iTUT and uTUT (see Table 3). This finding runs contrary to the prevailing notion of increased alpha band activity as a marker of mind-wandering and stands in contrast with most previous work employing tasks imposing demands on externally oriented attention (Arnau et al., 2020; Baldwin et al., 2017; Compton et al., 2019; Jin et al., 2019; Macdonald et al., 2011; Wamsley & Summer, 2020). However, studies using tasks relying on internally oriented attention, all reported reduced alpha activity during mind-wandering (Benedek, 2018; Braboszcz & Delorme, 2011; Rodriguez-Larios & Alaerts, 2021; van Son, De Blasio, et al., 2019; van Son, de Rover, et al., 2019), while increased alpha during tasks which do not require visual attention has been linked with improved processing of internal states (Cartocci et al., 2018). In a recent eye-tracking and EEG combined study, Ceh et al. (2020) observed a significant correlation between occipital alpha power and pupillary diameter during intertrial periods of rest, concluding that both are likely associated with a common gating mechanism in support of sustained internal attention with alpha increasing as a function of internal attention demands. This is in line with our results that intentional off-task thought is characterized by increased alpha measures, a pattern that is often observed when individuals are engaged in internal tasks (Benedek et al., 2014; N. R. Cooper et al., 2003; Hanslmayr et al., 2011; Katahira et al., 2018; Klimesch, 2012; Whitmarsh et al., 2014). Moreover, the patterns we observed in our data are broadly consistent with the hypothesis that iTUT and uTUT reflect differences in the role that top-down processes play in ongoing thought. More specifically, we hypothesize that iTUT is linked to the purposeful allotment of attention to internal processes and necessitates a neurophysiological mechanism for the top-down inhibition of irrelevant sensory input facilitated by alpha. Conversely, more unintentional TUTs are likely to result from intermittent failures to maintain attention on the task. A further indication supporting this interpretation is that the discriminative markers in the iTUT/uTUT contrast were similarly discriminative for the contrast between iTUT and the ‘distracted’ category of thought, while the contrast between uTUT and the ‘distracted’ category did not yield any significantly discriminative markers. Given that participants were instructed to report being distracted when they attended to the external environment besides the task, and that both states do not require active inhibition of sensory input, we speculate that the markers characterizing iTUT, e.g., alpha, reflect the application of control to organize an internal thought. In terms of markers characteristic of uTUT, delta normalized power was the most discriminative for this category compared to iTUT. Few studies have reported significant attentional state differences in the delta frequency range (for a review see J. W. Y. Kam et al., 2022) with one linking decreased frontal delta activity to off-task states when compared to on-task during SART (Wamsley & Summer, 2020). Notably, one recent study found that frontal delta activity predicted mind-wandering episodes (Andrillon et al., 2021). Given that slow cortical rhythms are typically characteristic of sleep, the increase of delta features we observed during uTUT may be indicative of a reduced alertness state. Interestingly, we also observed higher values of permutation entropy in the theta band as being discriminative of iTUT. Permutation entropy is an information complexity measure for time-series (Bandt & Pompe, 2002) which indexes the degree of conscious awareness in controls compared to anesthetized and minimally conscious state patients (Thul et al., 2016). Relatedly, Chen et al. (2020) used several different classifiers (e.g., SVM, random forest, naive Bayes, and k-nearest neighbors) with standard spectral measures as well as spectral entropy measures and found that the random forest classifier fared best with entropy-based features. Increased measures of complexity, including permutation entropy, have been suggested to be necessary for a specific representation to be selected for conscious processing (Sitt et al., 2014). Although we identified permutation entropy in the theta range as characteristic of iTUT, indicating increased conscious processing, the same measure of complexity in the beta and gamma range came out to be characteristic of uTUT, bearing out the relevance of such measures despite challenging interpretation.

**Table 3.** Summary of significant EEG markers from univariate analyses for all three classifications and MVPA accuracy for each classification. Markers are listed in descending order of AUC. Results shown for probe-caught (PC) vs. self-caught (SC) probe classification correspond to the under-sample method.

| Classification | On- vs. Off-Task | iTUT vs. uTUT |
| --- | --- | --- |
| **FDR corrected markers** | - | PE $\theta_{\text{mean}}$; $\alpha_{\text{mean}}$; $\vert \alpha\vert _{\text{mean}}$; $\beta_{\text{mean}}$; $\vert \delta\vert _{\text{mean}}$ |
| **Uncorrected markers** | $\vert \theta\vert _{\text{mean}}$; $\vert \theta\vert _{\text{std}}$; | $\alpha_{\text{std}}$; $\vert \alpha\vert _{\text{std}}$; PE $\beta_{\text{mean}}$; PE $\gamma_{\text{mean}}$ |
| **MVPA (AUC)** | 0.60 | 0.63 |

The second relevant finding is that theta features were characteristic of on-task states in contrast with off-task states. Our data suggest that periods reported as on-task are mainly characterized by increased normalized power at theta frequency, a pattern consistent across all comparisons. This agrees with previous findings of increased theta activity observed during executive functioning tasks and cognitive control (Cavanagh & Frank, 2014; Cavanagh & Shackman, 2015; P. S. Cooper et al., 2019). However, given discrepant findings about theta activity across studies using the SART - with some reporting increases and others decreases of theta activity during TUTs (see Kam et al., 2022 for a review) - and the small sample of on-task probe-caught reports in our study, it is difficult to draw any conclusive inference on the role of theta in the context of mind-wandering. Indeed, none of the discriminative markers for the contrast between on- and off-task survived correction for multiple comparison, most likely because of the low ratio of probe-caught reports to EEG markers. The iTUT/uTUT contrast did not suffer from the same issue and yielded multiple significant markers that survived correction, even though their discriminative power was comparable. Nevertheless, the AUC computation being less sensitive to sample size than *p* values, we suspect that the uncorrected significant markers presented here are reflective of underlying electrophysiological differences. Further demonstrating the discriminative ability of theta features, ERPs did not differentiate on-from off-task states, in opposition to our predictions based on prior studies that found reduced amplitudes of P1 and P3 components to be significant predictors of mind wandering (Dong et al., 2021; Groot et al., 2021; Martel et al., 2019). Nevertheless, our findings that oscillatory features were amongst the most discriminative for both contrasts are promising for the unobtrusive and continuous monitoring of EEG without the necessity for probing cognition to elicit and measure a brain response, as is the case for ERPs.

Our work could be affected by several limitations. The first one concerns the possibility of overfitting of our models in the on- and off-task contrast. This is a common issue with wide datasets containing many features and a small set of observations. Despite the assumption that intentional and unintentional TUTs have more in common than on- and off-task states, we obtained better classification performance for the former with a much larger sample size which significantly reduces the possibility of overfitting. Second, some of the differences reported might be attributable to the two types of thought probes used for both contrasts, namely probe-caught for the on-/off-task contrast and self-caught for the iTUT/uTUT contrast. Indeed, self-caught probes require participants to monitor their attention while performing the task which could have potentially introduced confounds. Third, although online thought sampling is a well-validated method to assess attentional states (Schubert et al., 2020; Welhaf et al., 2022), potential liability to biases (Seli, Jonker, et al., 2015; Weinstein et al., 2018) could have introduced noise and ultimately impacted classification accuracy. However, until robust, subject- and task-independent markers of attentional states are identified, self-reports remain indispensable. Lastly, although we were able to identify significant markers for the contrast between intentional and unintentional TUT, the discriminative markers for the on/off-task contrast did not survive correction for multiple comparisons, possibly due to the small sample size. Future work may include gathering more data points across different tasks to improve predictive performance and obtain task-independent EEG markers, with the goal of examining whether intentional and unintentional TUT can be predicted from ongoing EEG. Moreover, it is conceivable that the markers we obtained for attentional states are specific to the SART given that generalization of models typically leads to a drop in accuracy. With growing doubts concerning the utility of the SART in settling the ongoing debate surrounding the relationship between executive functions and mind-wandering (Boayue et al., 2020), future studies may want to extend our approach to paradigms with higher demands on executive resources such as the finger-tapping random generation task (Boayue et al., 2020; Groot et al., 2022). Future work may also profitably examine whether generating multi-modal representation by supplementing EEG measures with indirect markers of mind-wandering, e.g., behavioral or pupillometric measures, improves performance given the moderate success of these measures in previous work (Groot et al., 2021). Lastly, a key future direction concerns the modulation of TUT with brain stimulation to determine the causal relevance of cortical regions in mind-wandering. Selectively downregulating maladaptive types of mind-wandering, e.g., unintentional TUT, and upregulating advantageous types such as intentional TUT (Kam et al., 2022) may prove particularly beneficial. Our work represents a step forward in the direction of developing systems capable of detecting distinct attentional states in real-time and mitigating the negative effects of unintentional mind-wandering with promising applications in clinical and practical settings. Models providing reliable, real-time detection of TUTs, would be instrumental for the development of clinical interventions and real-world applications capable of mitigating the detrimental consequences of mind-wandering, as well as accelerate mind-wandering research by gradually replacing unreliable and disruptive, subjective measures with valid, objective measures.

To the best of our knowledge, ours is the first study to systematically contrast intentional and unintentional TUT by performing univariate and multivariate analyses on a broad and predefined set of EEG features, thus revealing distinct electrophysiological signatures characteristic for these two categories of thought. Intentional TUT was consistently disparate from other attentional states, as evidenced by significant differences across all comparisons in EEG markers linked to the top-down modulation of perception, e.g., increased alpha measures, presumably to shield internal processes by decoupling attention from sensory input. Together, our findings of unique electrophysiological signatures for conceptually distinct attentional states verifies the value of considering numerous EEG features, further substantiates the potential of EEG machine learning models for the detection of TUT, and adds to the ample evidence pointing to a distinction between intentional and unintentional mind-wandering (Banks & Welhaf, 2021; Golchert et al., 2017; Martínez-Pérez et al., 2021; Seli, Maillet, et al., 2017; Seli et al., 2016).

## Funding

This work was supported by a Marie Curie Fellowship (EU project 898813 CLONESA DLV 898813) awarded to Dr. Adrien Martel and by the ANR research grants (French National Research Agency) ‘OSCILOSCOPUS’ and Flag era JTC 2017 ‘CAUSALTOMICS’ awarded to Prof. Antoni Valero-Cabré.

## Author contribution

Conceptualization: AM, PMD, IHR

Data curation: AM

Investigation: AM, PMD

Methodology: AM, NB

Formal analysis: AM, NB

Supervision: AM, AVC, IHR, JDS, PMD

Visualization: AM, NB

Writing—original draft: AM, NB

Writing—review & editing: AM, AVC

## Supplementary Material

**Supplementary Figure 1.**
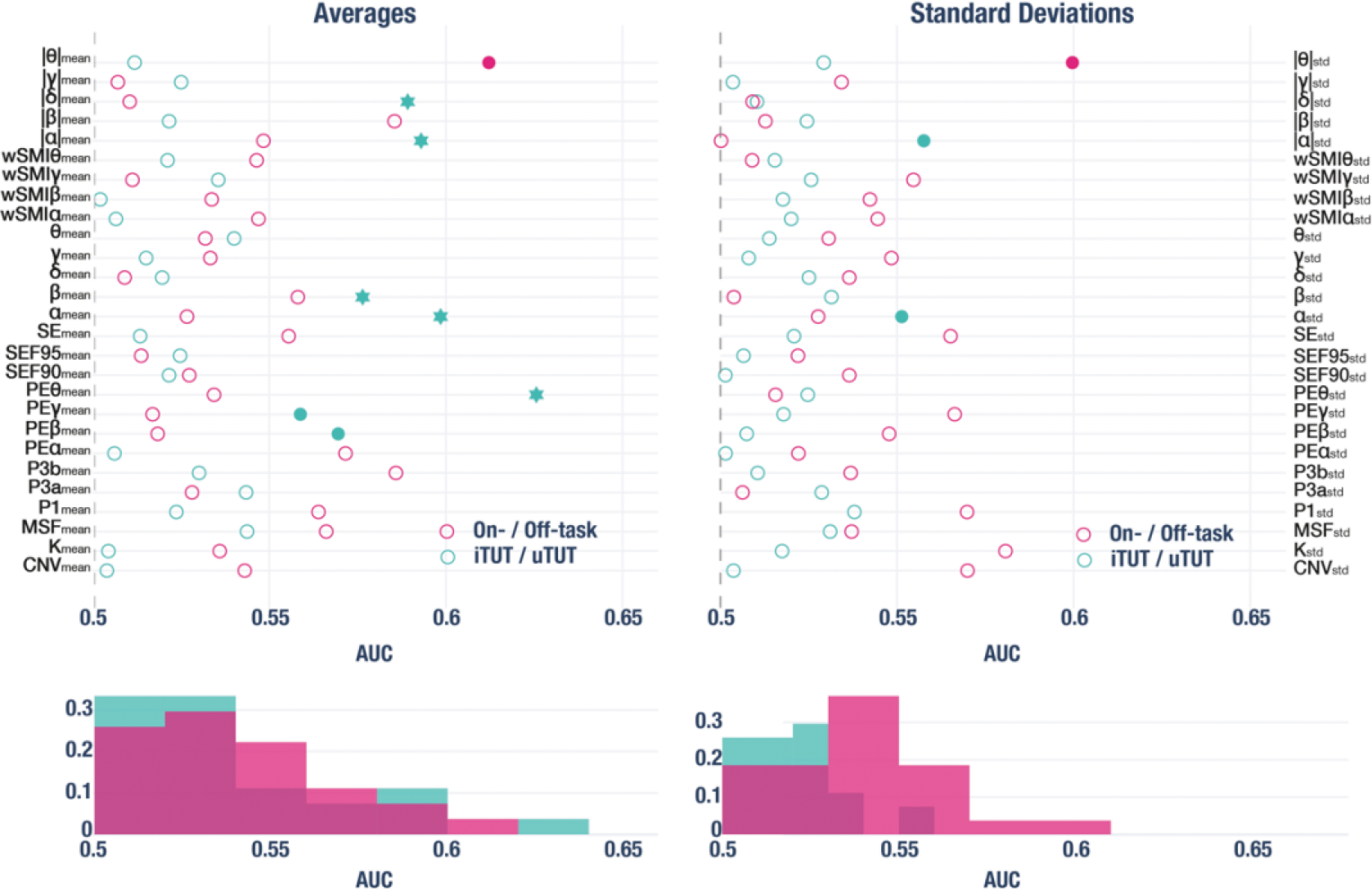
The mirrored area under the curve for all measures is the discrimination measure for the on-/off-task and iTUT/uTUT contrasts. All *AUC* < 0.50 were computed as the difference with 1 in order to mirror all the values. The markers were grouped together into averages (left) and standard deviations (right). The filled stars indicate *p* < 0.05 after FDR correction for multiple comparisons and the filled circle indicates *p* < 0.05 for the Mann Whitney U test before correcting for multiple comparisons. The lower histograms shows the same AUC distribution for each contrast for averages and standard deviations.

**Supplementary Figure 2.**
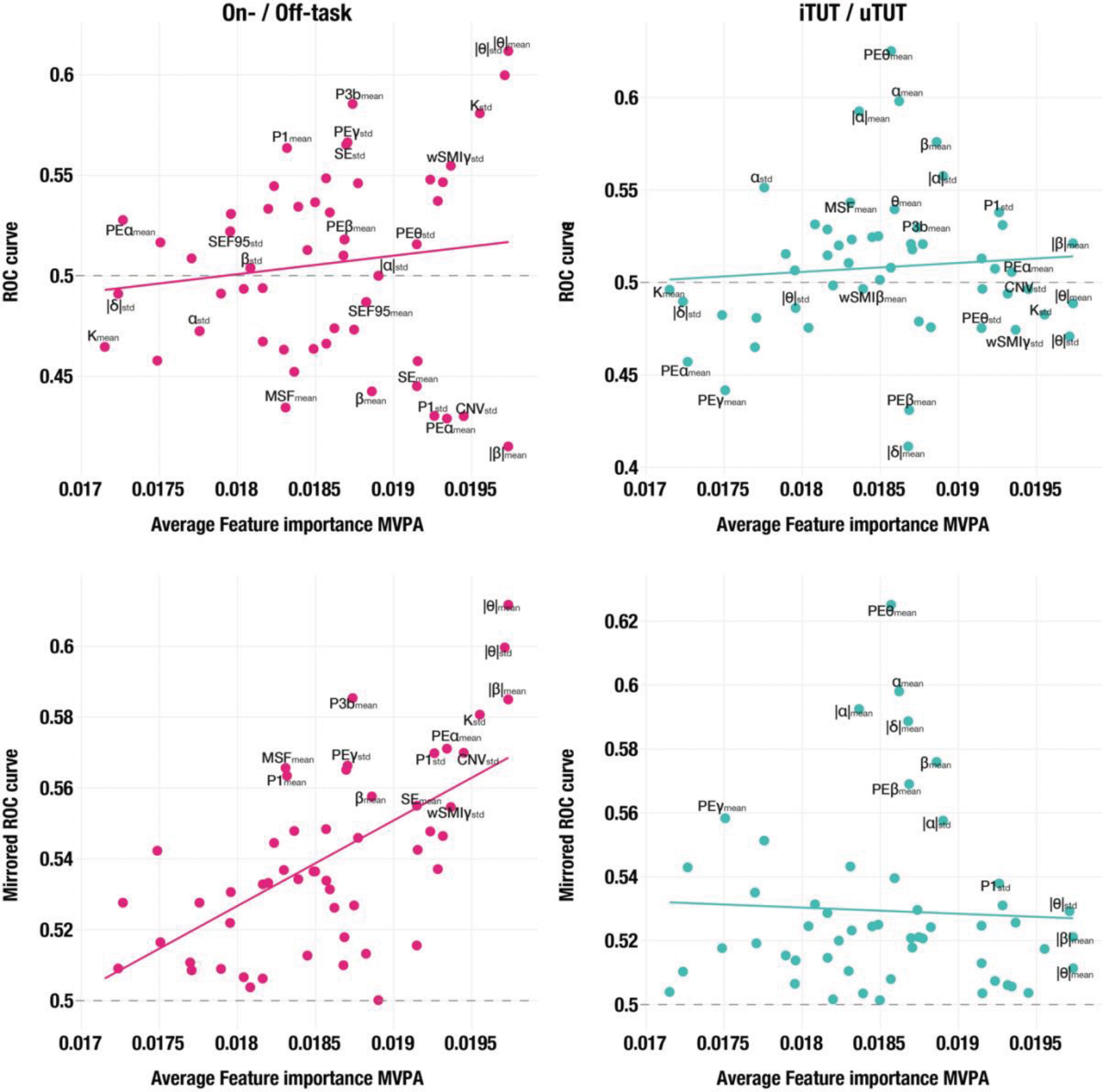
Comparison of univariate analysis and MVPA feature importance, with an ordinary least square line illustrating the trend between the two results. Upper quadrants display the comparison for on-/off-task (left) and iTUT/uTUT (right). Lower quadrants show the ROC univariate analysis with all AUC<0.50 values mirrored by calculating the difference from 1, aiding in visualizing potential trends.

**Supplementary Figure 3.**
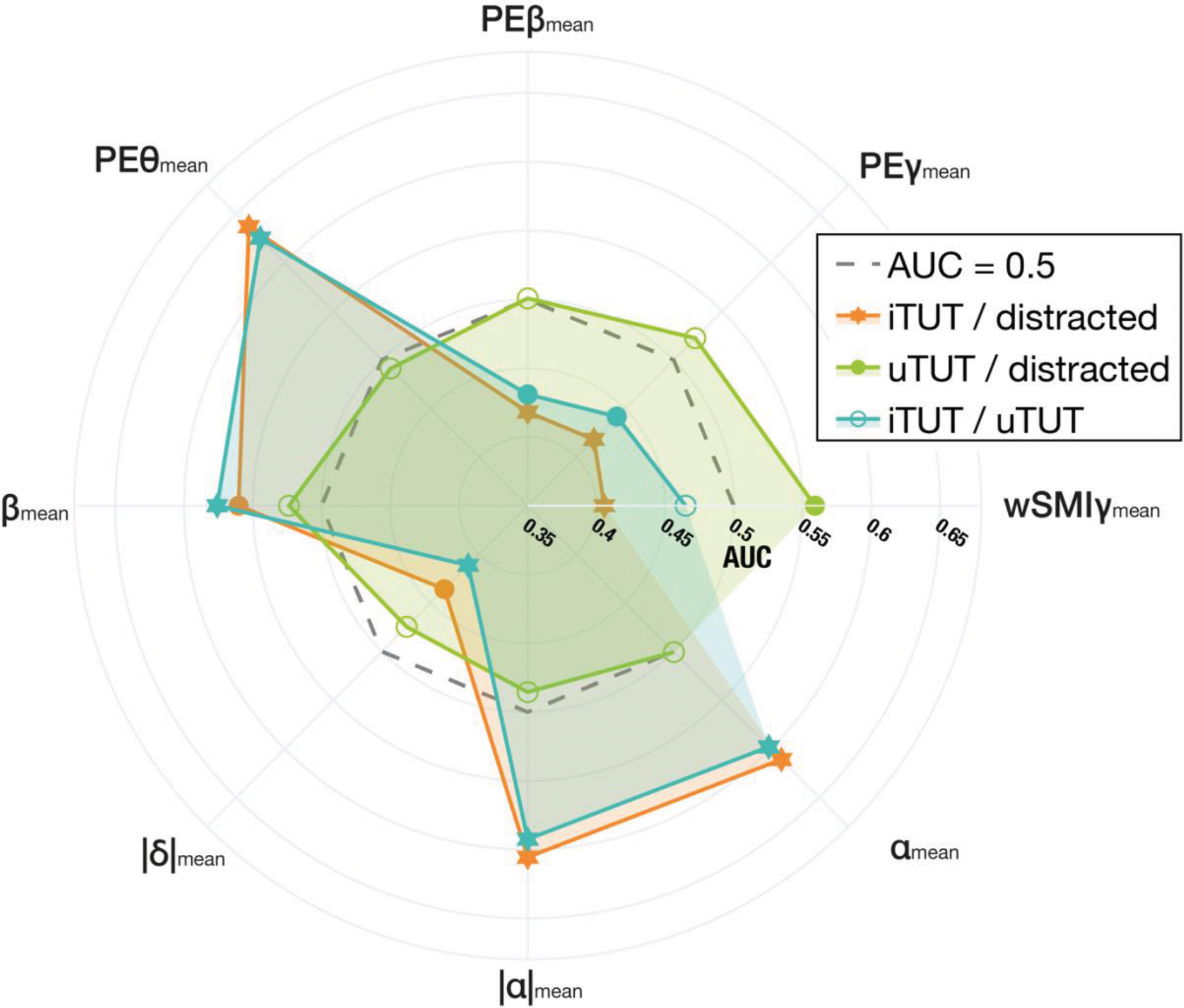
Scatter polar plot displaying all markers significant after multiple comparison correction for at least one of the iTUT/distracted, uTUT/distracted, and iTUT/uTUT comparisons. For the two comparisons involving iTUT, AUC>0.50 indicates a higher measure for iTUT than the other state; AUC<0.50 suggests the opposite pattern; and AUC=0.50 denotes no discrimination. For the uTUT vs. distracted comparison, AUC>0.50 indicates a higher measure for distracted than uTUT. Filled stars represent p<0.05 after FDR correction for multiple comparisons, and filled circles indicate p<0.05 for the Mann-Whitney U test before multiple comparison correction.

**Supplementary Table 1:**
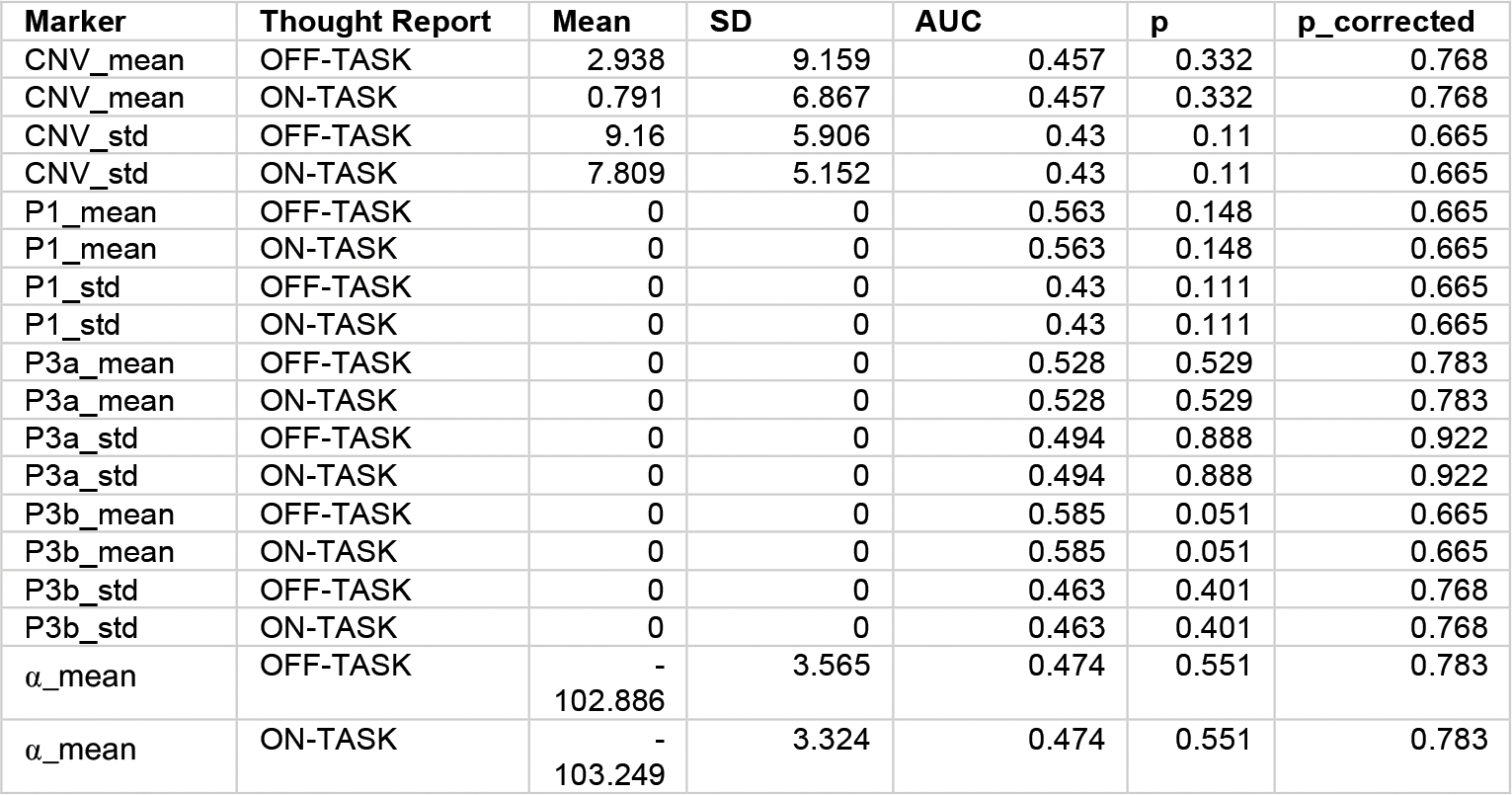
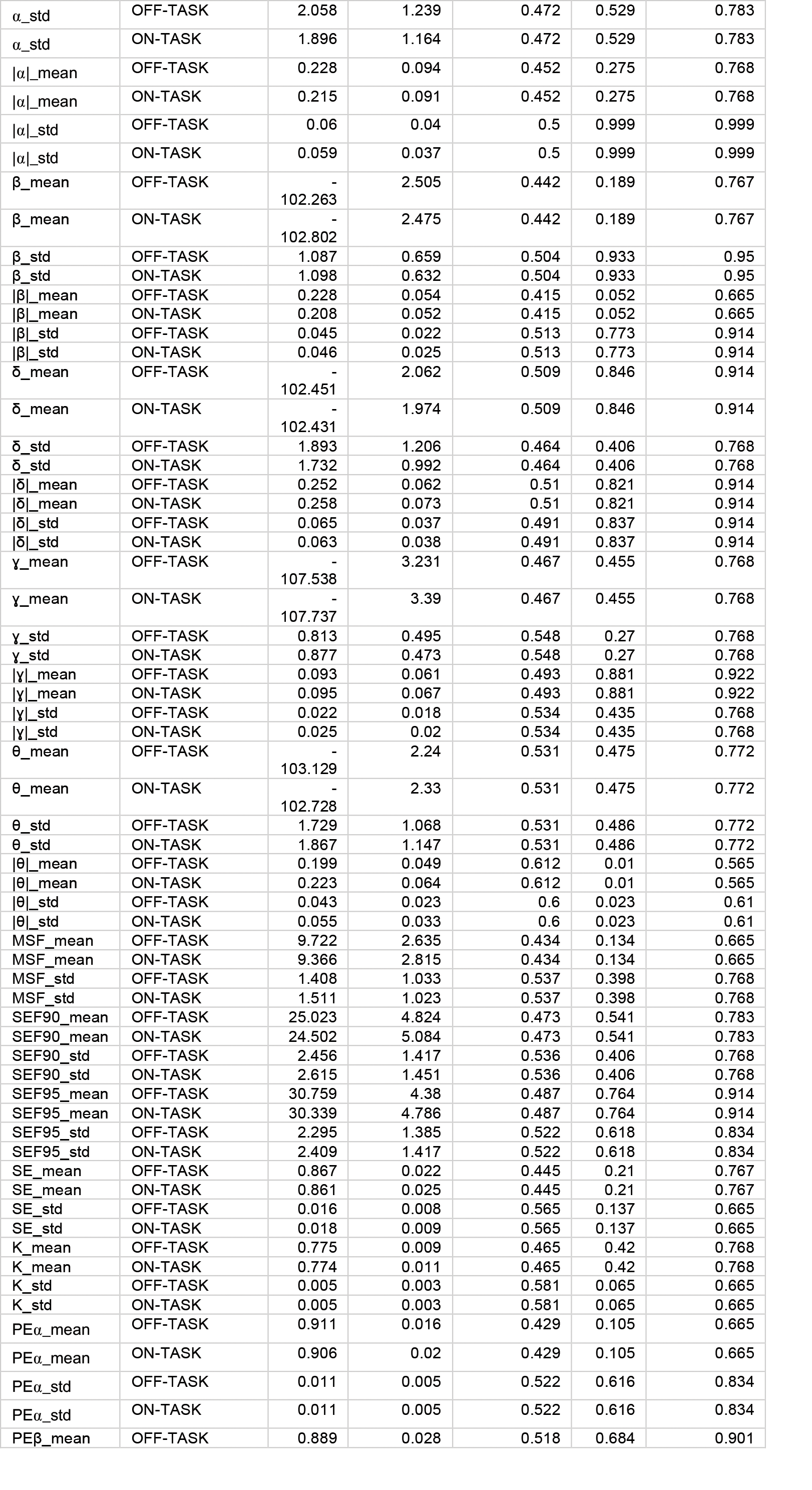
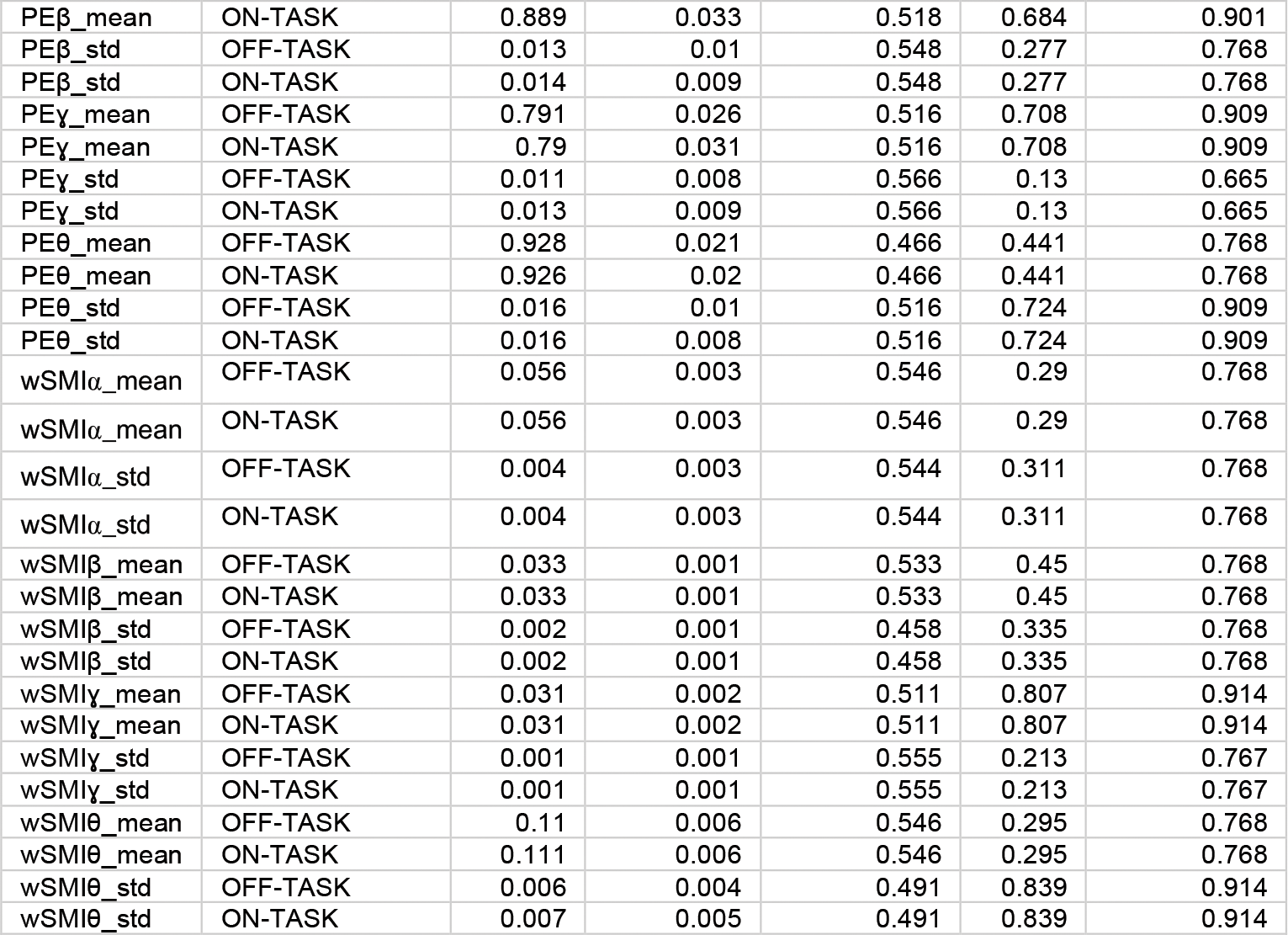
Comparison of On-Task and Off-Task Cognitive Markers: Means, Standard Deviations, AUC Values, and p Values (Uncorrected and Corrected)

**Supplementary Table 2:**
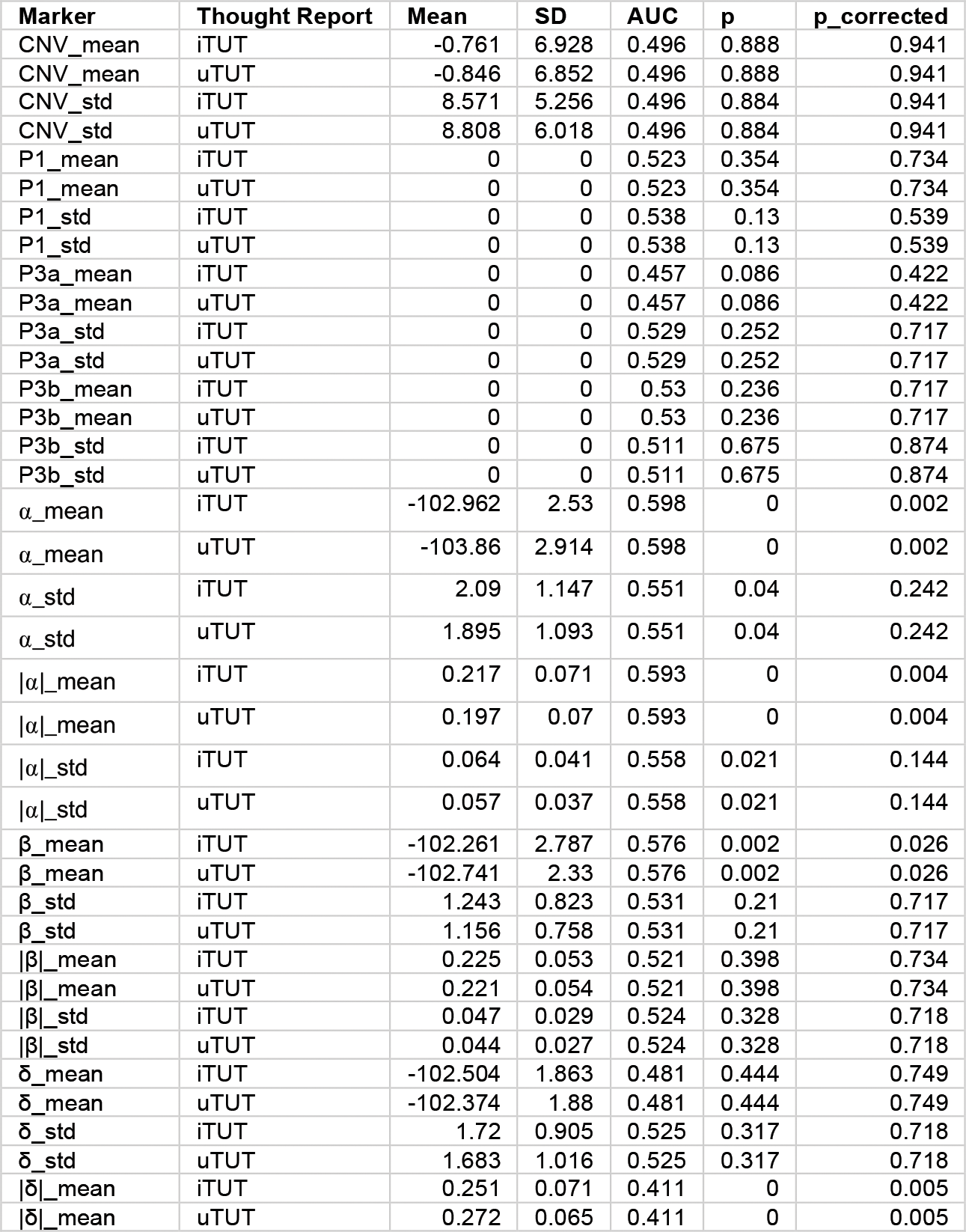
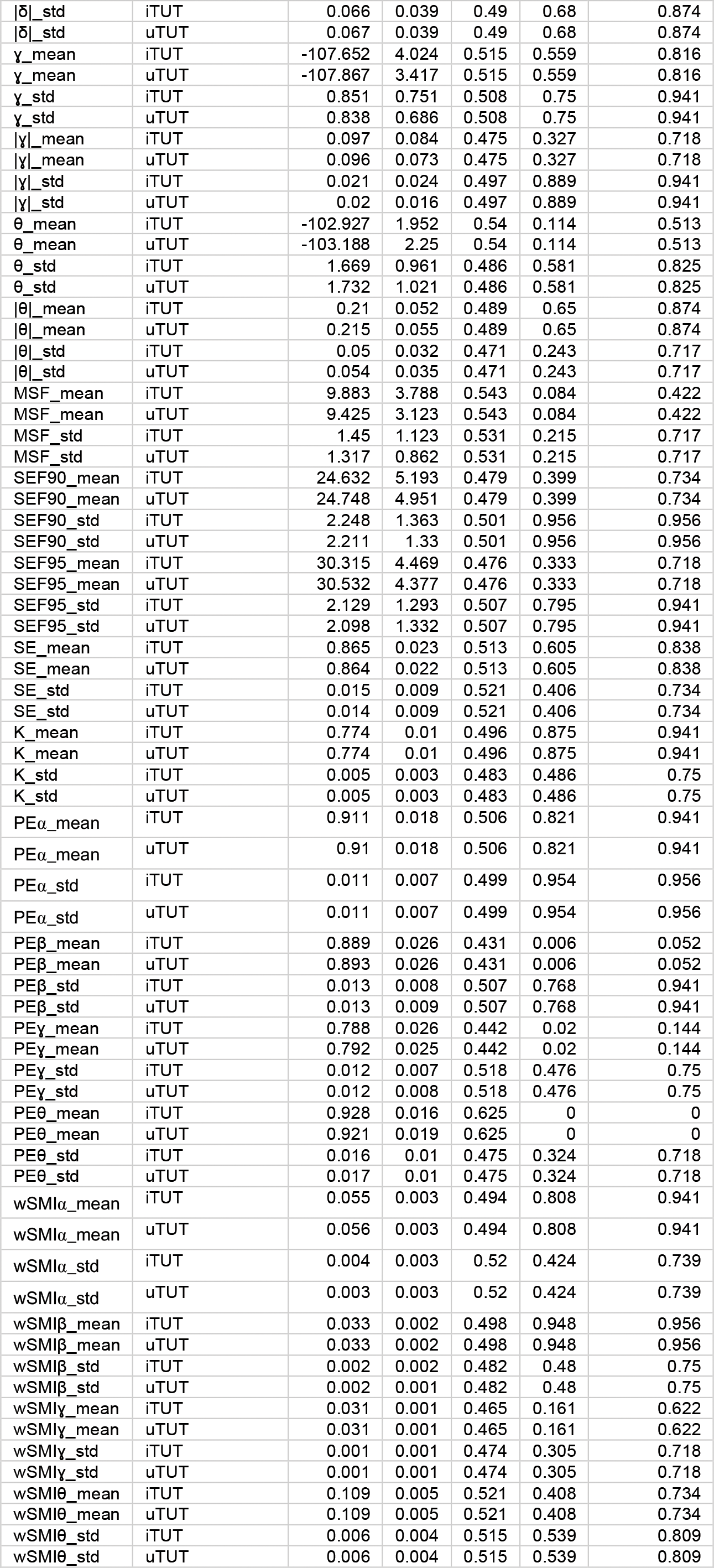
Comparative Analysis of Marker Values for iTUT and uTUT: Mean, Standard Deviation, AUC, and p Values (Uncorrected and Corrected).

| Marker | Thought Report | Mean | SD | AUC | p | p_corrected |
| --- | --- | --- | --- | --- | --- | --- |
| CNV_mean | iTUT | -0.761 | 6.928 | 0.496 | 0.888 | 0.941 |
| CNV_mean | uTUT | -0.846 | 6.852 | 0.496 | 0.888 | 0.941 |
| CNV_std | iTUT | 8.571 | 5.256 | 0.496 | 0.884 | 0.941 |
| CNV_std | uTUT | 8.808 | 6.018 | 0.496 | 0.884 | 0.941 |
| P1_mean | iTUT | 0 | 0 | 0.523 | 0.354 | 0.734 |
| P1_mean | uTUT | 0 | 0 | 0.523 | 0.354 | 0.734 |
| P1_std | iTUT | 0 | 0 | 0.538 | 0.13 | 0.539 |
| P1_std | uTUT | 0 | 0 | 0.538 | 0.13 | 0.539 |
| P3a_mean | iTUT | 0 | 0 | 0.457 | 0.086 | 0.422 |
| P3a_mean | uTUT | 0 | 0 | 0.457 | 0.086 | 0.422 |
| P3a_std | iTUT | 0 | 0 | 0.529 | 0.252 | 0.717 |
| P3a_std | uTUT | 0 | 0 | 0.529 | 0.252 | 0.717 |
| P3b_mean | iTUT | 0 | 0 | 0.53 | 0.236 | 0.717 |
| P3b_mean | uTUT | 0 | 0 | 0.53 | 0.236 | 0.717 |
| P3b_std | iTUT | 0 | 0 | 0.511 | 0.675 | 0.874 |
| P3b_std | uTUT | 0 | 0 | 0.511 | 0.675 | 0.874 |
| $\alpha$_mean | iTUT | -102.962 | 2.53 | 0.598 | 0 | 0.002 |
| $\alpha$_mean | uTUT | -103.86 | 2.914 | 0.598 | 0 | 0.002 |
| $\alpha$_std | iTUT | 2.09 | 1.147 | 0.551 | 0.04 | 0.242 |
| $\alpha$_std | uTUT | 1.895 | 1.093 | 0.551 | 0.04 | 0.242 |
| \|\mathbf{\alpha}\|_mean | iTUT | 0.217 | 0.071 | 0.593 | 0 | 0.004 |
| \|\mathbf{\alpha}\|_mean | uTUT | 0.197 | 0.07 | 0.593 | 0 | 0.004 |
| \|\mathbf{\alpha}\|_std | iTUT | 0.064 | 0.041 | 0.558 | 0.021 | 0.144 |
| \|\mathbf{\alpha}\|_std | uTUT | 0.057 | 0.037 | 0.558 | 0.021 | 0.144 |
| $\beta$_mean | iTUT | -102.261 | 2.787 | 0.576 | 0.002 | 0.026 |
| $\beta$_mean | uTUT | -102.741 | 2.33 | 0.576 | 0.002 | 0.026 |
| $\beta$_std | iTUT | 1.243 | 0.823 | 0.531 | 0.21 | 0.717 |
| $\beta$_std | uTUT | 1.156 | 0.758 | 0.531 | 0.21 | 0.717 |
| \|\mathbf{\beta}\|_mean | iTUT | 0.225 | 0.053 | 0.521 | 0.398 | 0.734 |
| \|\mathbf{\beta}\|_mean | uTUT | 0.221 | 0.054 | 0.521 | 0.398 | 0.734 |
| \|\mathbf{\beta}\|_std | iTUT | 0.047 | 0.029 | 0.524 | 0.328 | 0.718 |
| \|\mathbf{\beta}\|_std | uTUT | 0.044 | 0.027 | 0.524 | 0.328 | 0.718 |
| $\delta$_mean | iTUT | -102.504 | 1.863 | 0.481 | 0.444 | 0.749 |
| $\delta$_mean | uTUT | -102.374 | 1.88 | 0.481 | 0.444 | 0.749 |
| $\delta$_std | iTUT | 1.72 | 0.905 | 0.525 | 0.317 | 0.718 |
| $\delta$_std | uTUT | 1.683 | 1.016 | 0.525 | 0.317 | 0.718 |
| \|\mathbf{\delta}\|_mean | iTUT | 0.251 | 0.071 | 0.411 | 0 | 0.005 |
| \|\mathbf{\delta}\|_mean | uTUT | 0.272 | 0.065 | 0.411 | 0 | 0.005 |
| $\vert \delta\vert _{std}$ | iTUT | 0.066 | 0.039 | 0.49 | 0.68 | 0.874 |
| $\vert \delta\vert _{std}$ | uTUT | 0.067 | 0.039 | 0.49 | 0.68 | 0.874 |
| $\gamma_{mean}$ | iTUT | -107.652 | 4.024 | 0.515 | 0.559 | 0.816 |
| $\gamma_{mean}$ | uTUT | -107.867 | 3.417 | 0.515 | 0.559 | 0.816 |
| $\gamma_{std}$ | iTUT | 0.851 | 0.751 | 0.508 | 0.75 | 0.941 |
| $\gamma_{std}$ | uTUT | 0.838 | 0.686 | 0.508 | 0.75 | 0.941 |
| $\vert \gamma\vert _{mean}$ | iTUT | 0.097 | 0.084 | 0.475 | 0.327 | 0.718 |
| $\vert \gamma\vert _{mean}$ | uTUT | 0.096 | 0.073 | 0.475 | 0.327 | 0.718 |
| $\vert \gamma\vert _{std}$ | iTUT | 0.021 | 0.024 | 0.497 | 0.889 | 0.941 |
| $\vert \gamma\vert _{std}$ | uTUT | 0.02 | 0.016 | 0.497 | 0.889 | 0.941 |
| $\theta_{mean}$ | iTUT | -102.927 | 1.952 | 0.54 | 0.114 | 0.513 |
| $\theta_{mean}$ | uTUT | -103.188 | 2.25 | 0.54 | 0.114 | 0.513 |
| $\theta_{std}$ | iTUT | 1.669 | 0.961 | 0.486 | 0.581 | 0.825 |
| $\theta_{std}$ | uTUT | 1.732 | 1.021 | 0.486 | 0.581 | 0.825 |
| $\vert \theta\vert _{mean}$ | iTUT | 0.21 | 0.052 | 0.489 | 0.65 | 0.874 |
| $\vert \theta\vert _{mean}$ | uTUT | 0.215 | 0.055 | 0.489 | 0.65 | 0.874 |
| $\vert \theta\vert _{std}$ | iTUT | 0.05 | 0.032 | 0.471 | 0.243 | 0.717 |
| $\vert \theta\vert _{std}$ | uTUT | 0.054 | 0.035 | 0.471 | 0.243 | 0.717 |
| MSF_mean | iTUT | 9.883 | 3.788 | 0.543 | 0.084 | 0.422 |
| MSF_mean | uTUT | 9.425 | 3.123 | 0.543 | 0.084 | 0.422 |
| MSF_std | iTUT | 1.45 | 1.123 | 0.531 | 0.215 | 0.717 |
| MSF_std | uTUT | 1.317 | 0.862 | 0.531 | 0.215 | 0.717 |
| SEF90_mean | iTUT | 24.632 | 5.193 | 0.479 | 0.399 | 0.734 |
| SEF90_mean | uTUT | 24.748 | 4.951 | 0.479 | 0.399 | 0.734 |
| SEF90_std | iTUT | 2.248 | 1.363 | 0.501 | 0.956 | 0.956 |
| SEF90_std | uTUT | 2.211 | 1.33 | 0.501 | 0.956 | 0.956 |
| SEF95_mean | iTUT | 30.315 | 4.469 | 0.476 | 0.333 | 0.718 |
| SEF95_mean | uTUT | 30.532 | 4.377 | 0.476 | 0.333 | 0.718 |
| SEF95_std | iTUT | 2.129 | 1.293 | 0.507 | 0.795 | 0.941 |
| SEF95_std | uTUT | 2.098 | 1.332 | 0.507 | 0.795 | 0.941 |
| SE_mean | iTUT | 0.865 | 0.023 | 0.513 | 0.605 | 0.838 |
| SE_mean | uTUT | 0.864 | 0.022 | 0.513 | 0.605 | 0.838 |
| SE_std | iTUT | 0.015 | 0.009 | 0.521 | 0.406 | 0.734 |
| SE_std | uTUT | 0.014 | 0.009 | 0.521 | 0.406 | 0.734 |
| K_mean | iTUT | 0.774 | 0.01 | 0.496 | 0.875 | 0.941 |
| K_mean | uTUT | 0.774 | 0.01 | 0.496 | 0.875 | 0.941 |
| K_std | iTUT | 0.005 | 0.003 | 0.483 | 0.486 | 0.75 |
| K_std | uTUT | 0.005 | 0.003 | 0.483 | 0.486 | 0.75 |
| PEα_mean | iTUT | 0.911 | 0.018 | 0.506 | 0.821 | 0.941 |
| PEα_mean | uTUT | 0.91 | 0.018 | 0.506 | 0.821 | 0.941 |
| PEα_std | iTUT | 0.011 | 0.007 | 0.499 | 0.954 | 0.956 |
| PEα_std | uTUT | 0.011 | 0.007 | 0.499 | 0.954 | 0.956 |
| PEβ_mean | iTUT | 0.889 | 0.026 | 0.431 | 0.006 | 0.052 |
| PEβ_mean | uTUT | 0.893 | 0.026 | 0.431 | 0.006 | 0.052 |
| PEβ_std | iTUT | 0.013 | 0.008 | 0.507 | 0.768 | 0.941 |
| PEβ_std | uTUT | 0.013 | 0.009 | 0.507 | 0.768 | 0.941 |
| PEγ_mean | iTUT | 0.788 | 0.026 | 0.442 | 0.02 | 0.144 |
| PEγ_mean | uTUT | 0.792 | 0.025 | 0.442 | 0.02 | 0.144 |
| PEγ_std | iTUT | 0.012 | 0.007 | 0.518 | 0.476 | 0.75 |
| PEγ_std | uTUT | 0.012 | 0.008 | 0.518 | 0.476 | 0.75 |
| PEθ_mean | iTUT | 0.928 | 0.016 | 0.625 | 0 | 0 |
| PEθ_mean | uTUT | 0.921 | 0.019 | 0.625 | 0 | 0 |
| PEθ_std | iTUT | 0.016 | 0.01 | 0.475 | 0.324 | 0.718 |
| PEθ_std | uTUT | 0.017 | 0.01 | 0.475 | 0.324 | 0.718 |
| wSMIα_mean | iTUT | 0.055 | 0.003 | 0.494 | 0.808 | 0.941 |
| wSMIα_mean | uTUT | 0.056 | 0.003 | 0.494 | 0.808 | 0.941 |
| wSMIα_std | iTUT | 0.004 | 0.003 | 0.52 | 0.424 | 0.739 |
| wSMIα_std | uTUT | 0.003 | 0.003 | 0.52 | 0.424 | 0.739 |
| wSMIβ_mean | iTUT | 0.033 | 0.002 | 0.498 | 0.948 | 0.956 |
| wSMIβ_mean | uTUT | 0.033 | 0.002 | 0.498 | 0.948 | 0.956 |
| wSMIβ_std | iTUT | 0.002 | 0.002 | 0.482 | 0.48 | 0.75 |
| wSMIβ_std | uTUT | 0.002 | 0.001 | 0.482 | 0.48 | 0.75 |
| wSMIγ_mean | iTUT | 0.031 | 0.001 | 0.465 | 0.161 | 0.622 |
| wSMIγ_mean | uTUT | 0.031 | 0.001 | 0.465 | 0.161 | 0.622 |
| wSMIγ_std | iTUT | 0.001 | 0.001 | 0.474 | 0.305 | 0.718 |
| wSMIγ_std | uTUT | 0.001 | 0.001 | 0.474 | 0.305 | 0.718 |
| wSMIθ_mean | iTUT | 0.109 | 0.005 | 0.521 | 0.408 | 0.734 |
| wSMIθ_mean | uTUT | 0.109 | 0.005 | 0.521 | 0.408 | 0.734 |
| wSMIθ_std | iTUT | 0.006 | 0.004 | 0.515 | 0.539 | 0.809 |
| wSMIθ_std | uTUT | 0.006 | 0.004 | 0.515 | 0.539 | 0.809 |

## Notes

### Competing Interest Statement

The authors have declared no competing interest.

